# Multilevel computational approach to unlock the potential inhibitors of biofilm-EPS, persistence and quinolone signalling in *Pseudomonas aeruginosa* using mangrove-derived bioactive phytochemicals

**DOI:** 10.64898/2026.05.08.722855

**Authors:** Santosh Kumar Behera, Neelam Amit Kungwani, Yugal Kishore Mohanta

## Abstract

*Pseudomonas aeruginosa,* a Gram-negative opportunistic pathogen is well known for life-threatening acute infections among the human population. The bacterium can withstand most antibiotics by using their high levels of inherent and acquired resistance mechanisms such as Biofilm-EPS, Persistence, and Quorum sensing (QS). Owing to the importance of adaptive antibiotic multi-drug resistance of *P. aeruginosa,* the current investigation is aimed to explore the phytochemicals derived from mangrove plants as potential agents to control biofilm and drug resistance mechanisms through a multi-mechanistic computational approach. For identifying potential compounds and target, *In-silico* drug repurposing technique is implemented by docking/virtual screening of 49 phytochemical compounds against 18 proteins involved in the Persister Cell formation, QS, and EPS synthesis in *P. aeruginosa* which resulted the proteins RelA and SpoT (persistence), PqsA, and PqSR (QS), and PelA and PelB (EPS synthesis) and compounds Taraxerone and Taraxerol to be potential. The results of docking were well corroborated with MD simulations. These targets and compounds explored through *in-silico* approach, are found to target potential antimicrobial pathways involving EPS synthesis, persistence genes, and QS, aiming to enhance antibiotic efficacy. Further, this study could be reference for *in-vivo* and *in-vitro* investigations to evaluate the further effectiveness of the compounds and potentiality of the proteins for MDR therapeutics of *P. aeruginosa*.

## 1. Introduction

*Pseudomonas aeruginosa* is an opportunistic pathogen that causes a wide spectrum of infections in humans and animals. In humans, the bacterium causes array of infections both community- and hospital-acquired. The bacterium is also associated with infection of the skin, eyes, urinary tract and ears. Additionally, it causes diseases in livestock and companion animals, including otitis and urinary tract infections in dogs, mastitis, and endometritis in dairy cows [1]. Infections caused by *P. aeruginosa* are linked to its biofilm formation and quorum sensing (QS) that have been studied in diverse host ranges. Biofilm is a surface-attached growth where cells are enclosed in a matrix of extracellular polymeric substances (EPS) where cell-cell communication is synchronized by QS. The bacterium produces several EPS such as alginate, Pel, PsI and PelD crucial for its biofilm development. Biofilm-EPS inactivates antibiotics by conferring impermeability, quenching activity, and enzymatic degradation. The Las, Rhl and PQS are three major QS systems in *P. aeruginosa* which regulate hundreds of genes. The Lux-like AHL synthase LasI and RhlI regulate several genes associated with biofilm development and pathogenesis. The PQS on the other side involved in *P. aeruginosa* virulence including antibiotic-induced persistence [2], [3], [4].

Persister cells (PC) are phenotypically antibiotic-tolerant and have been linked to antibiotic treatment failures. They are metabolically inactive cells commonly reported under antibiotic stress, nutrient deprivation, and biofilms. The majority of the PC studies have been focused on *E. coli* and *Staphylococcus aureus*, but *P. aeruginosa* has not been explored widely. In recent years a number of genes *spoT, relA, dksA, spuC, ycgM, dinG, pilH*, Toxin-Antitoxin (TA) modules have been reported to be associated with PC formation [5], [6].

Since monotherapy cannot eradicate PC and biofilms, combinational antibiotic therapies are employed to control such types of infection, which may result in multidrug resistance (MDR). To combat antimicrobial resistance (AMR) spread, phytochemicals are potential alternatives to antibiotics which can control biofilm-related infections. Alternatively, phytochemicals can be used as an antibiotic adjuvant to potentiate drug efficiency against infection-causing organisms [7], [8]. Plants are a unique source of antimicrobial compounds and are popularly used for the treatment of bacterial infections under the herbal medication system. Similar to terrestrial plants, mangrove plants are also rich in therapeutic phytochemicals. Mangroves are salt-tolerant plants grown predominantly in intertidal regions. The therapeutic knowledge of mangroves came from coastal and traditional communities and are the source of unique bioactive phytochemicals, with diverse pharmacological properties [9], [10]. Mangroves and their associates have been reported as a source of new eras of secondary metabolites and phytochemicals. *Sonneratia apetala, Rhizophora mucronata, Avicennia marina, Avicennia officinalis Lumnitzera racemosa, Excoecaria agallocha* etc. are known to possess antibacterial properties. Several promising but underutilized phytochemicals found in mangrove plants can be used to manage bacterial biofilms and associated antibiotic resistance [10], [11], [12], [13].

In this study, a multilevel computational approach targeted 49 phytochemicals derived from *Avicennia officinalis* and *Avicennia marina* against 18 protein targets of *P. aeruginosa* that cause multiple potential infectious effects. The majority of the computational approaches on phytochemical repurposing only focus on the RhlI and LasI QS networks. In this very first study, phytochemicals were identified with anti-biofilm-EPS, anti-persistence, and anti-quorum sensing properties against the potential antimicrobial targets with EPS synthesis, persistent genes, along with QS to potentiate antibiotic activities.

## 2. Materials and Methods

### 2.1. Sequence, structure, and functional analysis target proteins

For the analysis of the sequence, structure, and function of 18 proteins involved in the PC formation, QS, and EPS synthesis in *P. aeruginosa,* information was retrieved from the UniProtKB database (https://www.uniprot.org/) utilizing various protein IDs and related literature. The proteins’ experimental configurations were sourced from the Protein Data Bank (PDB). Since specific proteins or targets are not made accessible or the 3D experimental structures in PDB are not suitable, a theoretical 3D structure has been opted for from the alpha fold protein structure database, which EBI has anticipated and tracked up to date.

### 2.2. Prediction of binding site

Web servers namely Computed Atlas of Surface Topography of Proteins (CASTp) http://cast.engr.uic.edu, DEPTH (http://cospi.iiserpune.ac.in/depth), Grid-based Hemi pocket finder (GHECOM) (https://pdbj.org/ghecom/) and PDB sum were employed for the prediction of the binding sites of the targeted proteins using the adhesion execution of the servers [14]. Leveraging the AutoDock 4.2 tool, the targeted protein’s coordinates (x, y, and z) were assigned [15].

### 2.3. Mining of phytochemicals and virtual screening

An array of databases like PubMed, Ovid Scopus, and academic search engines were used for the investigation of the literature pertinent to phytochemicals derived from *Avicennia* genus and reported for antimicrobial/antibiofilm activity against bacterial pathogens. Plants from this genus are rich in flavonoids, naphthalene derivatives, terpenoids, and glycosides with therapeutic effects [10]. Stem, root and leaf extract of *A. marina* and *A. officinalis* have been reported for antibacterial and antibiofilm activity against various ESKAPE pathogens [10], [16], [17], [18].

The structural information of 49 phytochemical compounds (PCC) **(Table 1)** was obtained from the PubChem database with various Compound ID’s in Structure Data Format (SDF). For the structural non-availability of the PCC’s, ChemDraw20.0 was used to draw in .cdx format, subsequently, the 3D structure of the PCC’s was converted to .pdb format using Chem3D 20.0 and BIOVIA Discovery Studio 4.5 Visualizer (BIOVIA, San Diego, CA, USA) which is mostly preferred format for different docking tools. All these 49 PCC were subjected to virtual screening against 18 identified corresponding targeted proteins through Perl script using Autodock Vina platform[19].

**Table 1:**
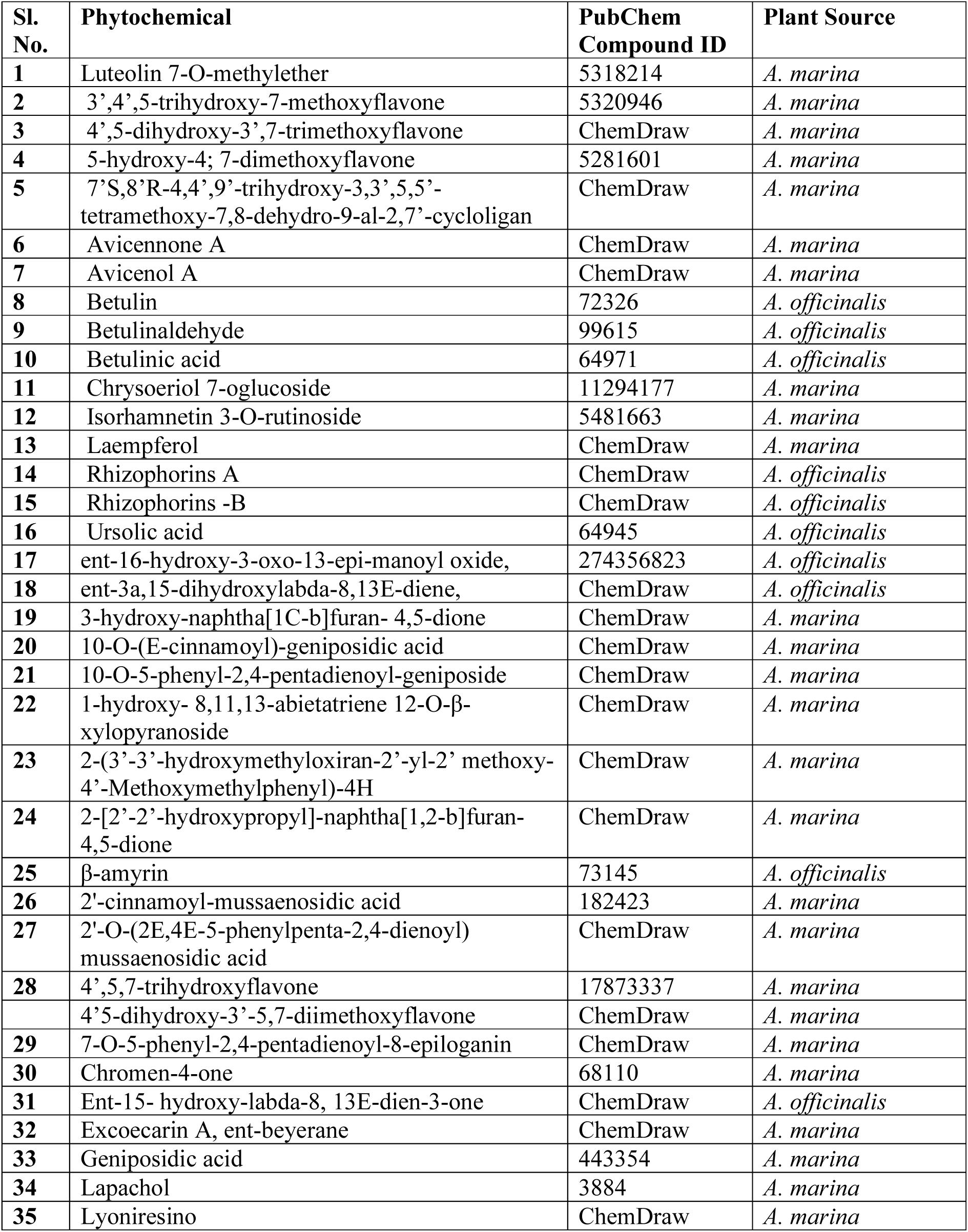
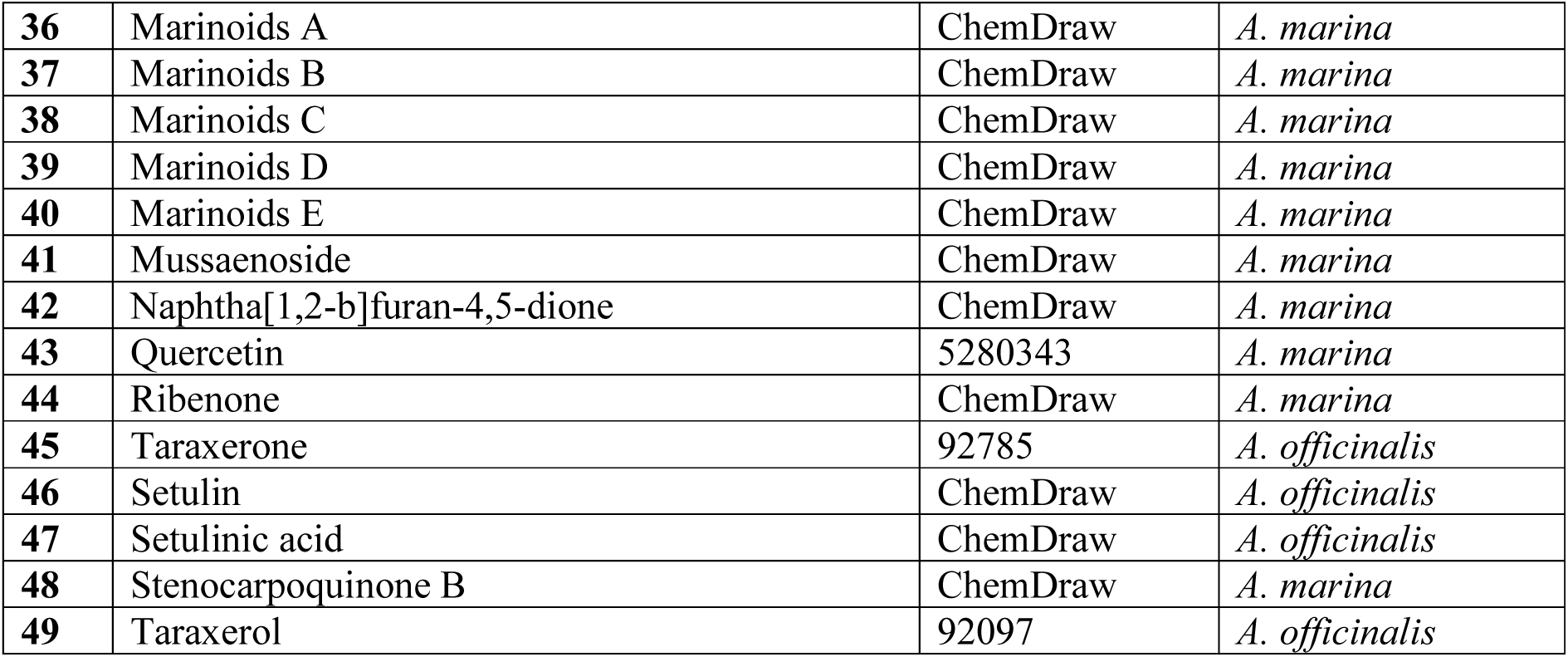
List of phytochemicals virtual screened against 18 proteins involved in the PC formation, QS, and EPS synthesis in *P. aeruginosa*.

### 2.4. Molecular Docking

Glide (Grid-based Ligand Docking with Energetics), Schrödinger, LLC, New York, NY, 2022 (Schrödinger Release 2022-4: Maestro, Schrödinger, LLC, New York, NY, 2022) was employed for docking analysis of top-ranked compounds resulted from virtual screening against the proteins associated with persistence, QS, and EPS synthesis in *P. aeruginosa* in extra precision (XP) mode. According to binding energy values, intermolecular hydrogen (H)-bonds, and other hydrophobic and electrostatic interactions, the most promising docked complexes have been classified and assembled for a subsequent computational study. The Schrödinger ligand interactions module was deployed to establish the potential existence of intermolecular links in protein-ligand complexes.

### 2.5. Molecular dynamics (MD) simulations

When it comes to investigating and projecting atom and molecule dynamics underneath the constraints of macromolecular structure-to-function acquaintances, molecular dynamics (MD) is a prominent computational approach [20], [21]. The system’s dynamic “ evolution “ becomes apparent by allowing the atoms and molecules to interact for a specified time [22]. It offers instantaneous scrutiny of the molecular mechanisms enabling aptamer-receptor binding and renders optimum receptor binding scenarios for theranostic alternatives [23]. To verify drug binding modalities as well as offer an extensive understanding of the protein-ligand complex, we executed MD simulations of the Apo (only protein) and Holo (protein-ligand complex) using the Desmond module of Schrödinger, LLC, New York, NY, 2022 (Schrödinger Release 2022-4: Maestro, Schrödinger, LLC, New York, NY, 2022). An MD simulation spanning 100 nanoseconds (ns) is carried out on the ligand-protein assemblages with the highest scores. Minimization, heating, equilibration, and manufacturing were the steps in the MD protocol. The MD process comprises minimization, heating, equilibration, and the fabrication run [24]. Protein-ligand complexes have been optimized leveraging the OPLS4 force field, and atomic coordinates and topology were automatically defined [25]. The compound became immersed in an orthorhombic box (15 x 15 × 10 Å), adopting the SPC solvent model. The process of neutralization of the physiological pH was succeeded by adding 0.15 M NaCl. The water box was configured with the Particle Mesh Ewald (PME) boundary condition to ensure no solute atoms occurred beneath 10 Å of the border. A combination of the NPT ensemble, the whole system was simulated at 300 K for 100 ns. Using root mean square deviation (RMSD) and root mean square fluctuation (RMSF) plots, the protein’s structure and dynamic movement were analysed. RMSD assesses the modification in a protein’s backbones between its original structural conformation and ultimate position. A protein or complex’s flexible region can be positioned using the RMSF proximity [26]. The simulation interaction schematic illustrates the most probable ligand binding mode at the protein binding site [27].

## 3. Results

### 3.1. Sequence, structural information of screened target proteins

Taking together the existing reports and the results of virtual screening it was depicted that the genes *relA* and *spoT* and their corresponding proteins are associated with persistence, *pqsA*, and *pqSR* with quorum sensing, and *pelA* and *pelB* with EPS synthesis in *P. aeruginosa.* The detailed information of RelA and SpoT protein targets was sourced from UniProtKB database with ID: A0A072ZLF8 (A0A072ZLF8_PSEAI) and A0A069Q2V6 (A0A069Q2V6_PSEAI) respectively. The 3D structure RelA and SpoT targets were retrieved from the alpha fold protein structure database. Similarly, the sequence, structure, and functional information of PqsA, and PqsR was retrieved from UniProtKB database with ID: Q9I4X3 (PQSA_PSEAE) and A0A7G9KWC2 (A0A7G9KWC2_PSEAI) respectively.

*Pseudomonas aeruginosa’*s PqsA X-ray diffraction structure (PDB ID: 5OE3), which has an estimated length of 398 (2-399) amino acids (aa), was compiled from the PDB database with a spatial resolution of 1.43Å. The 3D structure PqsR was retrieved from the alpha fold protein structure database. Subsequently, the 3D structures of PelA and PelB were retrieved from PDB database with PDB ID’s- 5TCB and 5WFT with resolutions 1.53Å and 2.82Å.

### 3.2. Annotation of the binding sites of the targeted proteins

All web servers’ consensus assessments identified substantial residues from their specific protein targets that contribute to the proliferation of active sites. Using ADT v.1.5, Kollman charges were attributed to each of the 18 protein targets, and then compounds were entrusted Gasteiger partial charges. The amino acid residues of the respective screened proteins that actively participate in active site/binding pocket formation were used for grid generation with specified dimensions to facilitate the ligand/compounds’ fully flourished shape, as apparent in **Table 2**.

**Table 2.** Details of the grid dimensions of respective reported protein targets that contribute in binding pocket formation.

| Sl. No. | Name of the proteins | Grid dimensions |
| --- | --- | --- |
| 1. | RelA | x-centering: -12.096,<br>y-centering: -42.797,<br>z-centering: -17.25 |
| 2. | SpoT | x-centering: -6.763,<br>y-centering: -19.471,<br>z-centering: -5.447 |
| 3. | PqsA | x-centering: -6.8,<br>y-centering: 1.118,<br>z-centering: 5.876 |
| 4. | PqSR | x-centering: -6.927,<br>y-centering: 7.005,<br>z-centering: 19.608 |
| 5. | PelA | x-centering: 43.172,<br>y-centering: 25.099,<br>z-centering: 15.589 |
| 6. | PelB | x-centering: 6.247,<br>y-centering: -20.613,<br>z-centering: -122.7 |

### 3.3. Screening of phytochemical compound against the targeted proteins

In the present investigation, 49 PCC were screened against the 18 proteins associated with persistence, QS, and EPS synthesis in *P. aeruginosa.* The virtual screening results depicted the six potential protein targets out of 18 with persistence (RelA and SpoT), QS (PqsA, and PqSR), and EPS synthesis (PelA and PelB) through Perl script using Autodock Vina platform. The virtual screening resulted in the compounds Taraxerone and Taraxerol better interaction with almost all the above-mentioned targets on the basis of their binding energy which was further examined for its potential activity (**Supplementary Table 1**). Between, the two compounds, Taraxerone depicted the binding energy of -7.8 & -8.1 kcal/mol against RelA and SpoT, and -7.7 & -8.3 kacl for Taraxerol; -10.5 and -10.3 kcal/mol against PqsA, and PqsR in Taraxerone and -10.2 and -9.5 kcal/mol in case of Taraxerol. Subsequently, -8.5 and -9.4 kcal/mol against PqsA, and PqsR in Taraxerone and -9.0 and -8.1 kcal/mol in the case of Taraxerol. Molecular docking was further hypothesized based on the binding energy and substantiated by utilizing various algorithms for computation.

### 3.4. Molecular docking

The ligand shape and perspective extrapolation into a predetermined binding site, also referred to as an active site, is a significant period in the docking process. Considering the consensus results of virtual screening the activity of compounds, docking of Taraxerone and Taraxerol against the different protein targets was performed using Glide, Schrödinger, LLC, New York, NY, 2022 (Schrödinger Release 2022-4: Maestro, Schrödinger, LLC, New York, NY, 2022) module in extra precision (XP) mode. Ten conformations of each protein-ligand combination received collection from the docking assays; a particularly promising posture defined by intermolecular H-bonds and binding energy is preserved for additional investigations.

The binding energies and other interaction studies of all the twelve complexes i.e, Holo1: RelA- Taraxerone complex, Holo2: RelA - Taraxerol complex, Holo3: SpoT - Taraxerone complex, Holo4: SpoT - Taraxerol complex, Holo5: PqsA - Taraxerone complex, Holo6: PqsA – Taraxerol complex, Holo7: PqsR- Taraxerone complex, Holo8: PqsR-Taraxerol complex, Holo9: PelA- Taraxerone complex, Holo10: PelA -Taraxerol complex, Holo11: PelB- Taraxerone complex, Holo12: PelB-Taraxerol complex, is presented in Table 3, Fig. 1A-1L. The drug-target complexes’ binding frequencies varied, relying on the docking data. Only the most opportune position with the highest binding energy was meticulously chosen for the intermolecular interaction analysis, which consisted of several conformations conserved from the docking research analysis utilizing the GLIDE module. The docking study reveals that the docking energies -3.904 kcal/mol, for PqsA – Taraxerol complex, and - 2.211 kcal/mol for RelA- Taraxerone complex complex, as the highest and lowest among all the docked protein-ligand complexes. The 12 docked complexes underwent MD simulations intending to clarify and corroborate the binding dynamics of the protein-ligand interaction during an allocated frame.

**Figure 1:**
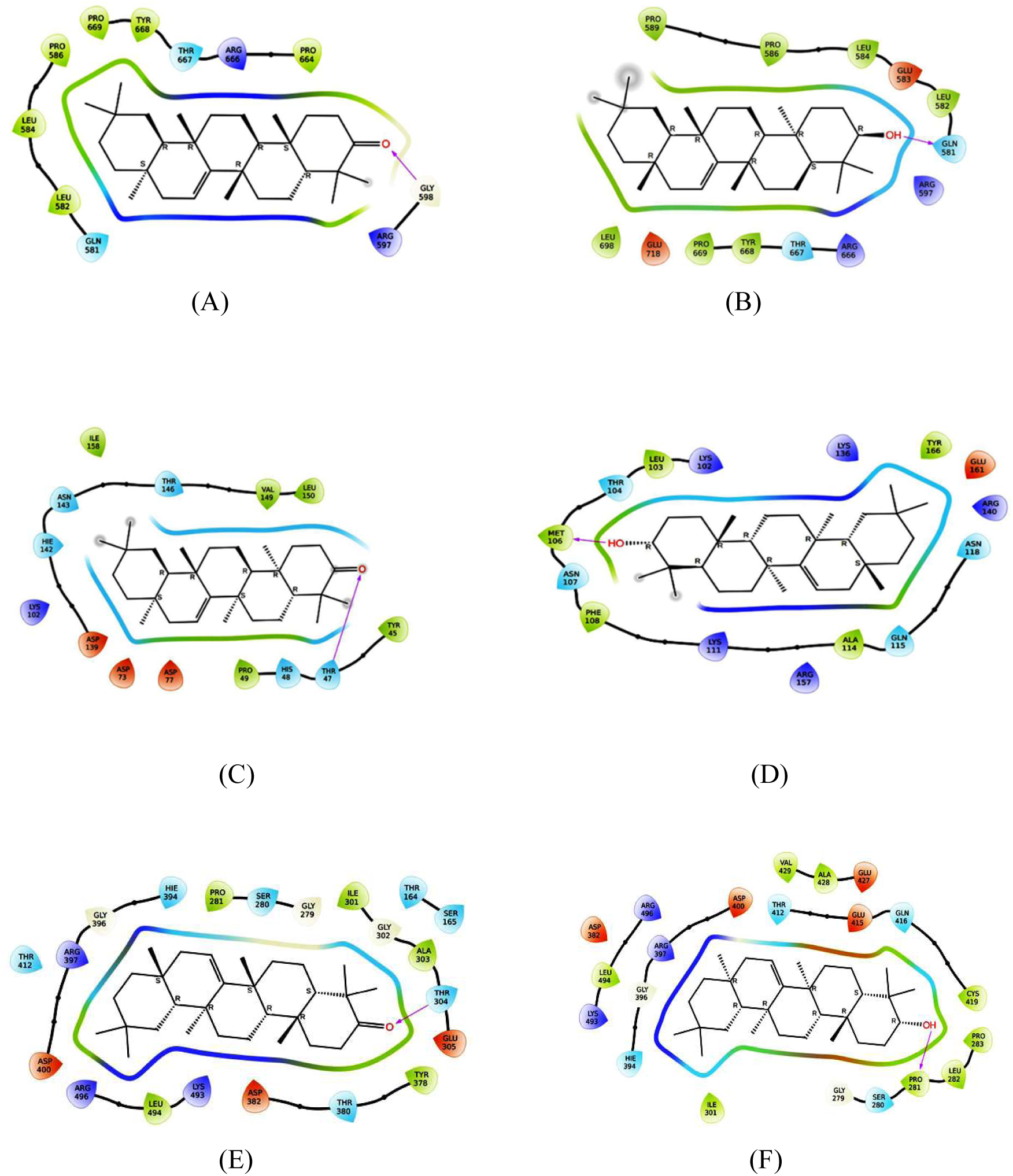
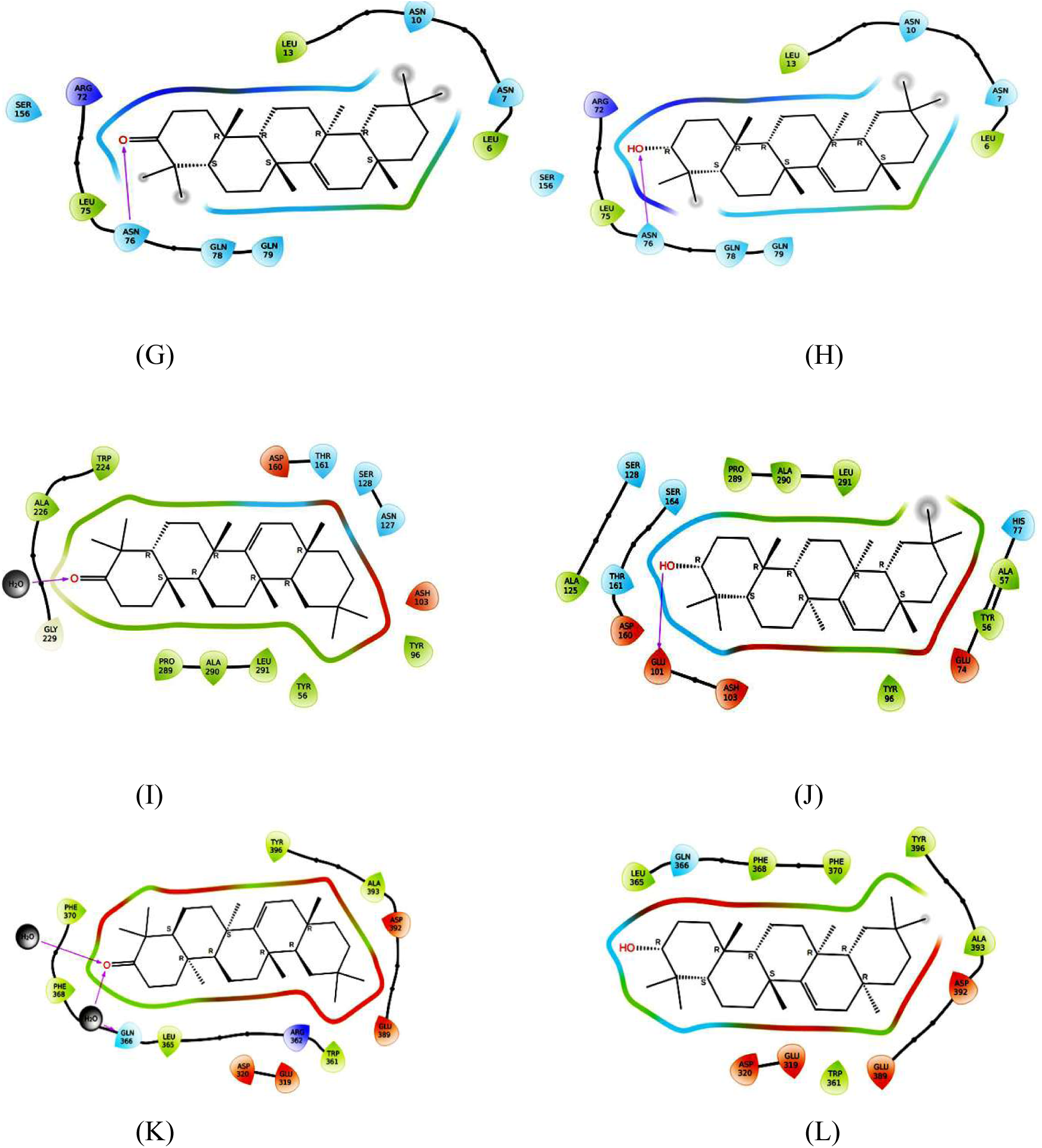
Intermolecular hydrogen bonding, electrostatic and hydrophobic interactions formed between various proteins and compounds Taraxerone and Taraxerol (**A)** RelA-Taraxerone complex, **(B)** RelA - Taraxerol complex, **(C)** SpoT - Taraxerone complex, **(D)** SpoT - Taraxerol complex, **(E)** PqsA - Taraxerone complex, **(F)** PqsA – Taraxerol complex, **(G)** PqsR- Taraxerone complex, **(H)** PqsR-Taraxerol complex, **(I)** PelA- Taraxerone complex, **(J)** PelA -Taraxerol complex, (K) PelB- Taraxerone complex, **(L)** PelB-Taraxerol complex. The images were obtained by ligand interactions module of Schrödinger.

**Table 3:** Molecular docking analysis of 2 compounds against six different proteins associated with persistence, Quorum Sensing and EPS synthesis.

| **Sl. No.** | **Target** | **Compound** | **Binding Energy(kcal/mol)** | **No. of H - Bonds** | **H-Bond Forming Residues** | **Average Distance of H-Bonds (Å)** |
| --- | --- | --- | --- | --- | --- | --- |
| **1.** | RelA | Taraxerone | -2.211 | 1 | Gly598 | ~ 1.929 |
| **2.** | RelA | Taraxerol | -2.732 | 1 | Gly581 | ~ 1.840 |
| **3.** | SpoT | Taraxerone | -2.285 | 1 | Thr47 | ~ 2.106 |
| **4.** | SpoT | Taraxerol | -3.102 | 1 | Met106 | ~2.026 |
| **5.** | PqsA | Taraxerone | -3.626 | 1 | Thr304 | ~1.995 |
| **6.** | PqsA | Taraxerol | -3.904 | 1 | Pro281 | ~2.557 |
| **7.** | PqsR | Taraxerone | -3.163 | 1 | Asn76 | ~2.115 |
| **8.** | PqsR | Taraxerol | -3.289 | 1 | Asn76 | ~2.073 |
| **9.** | PelA | Taraxerone | -2.541 | 1 | Tpr224 | ~1.916 |
| **10.** | PelA | Taraxerol | -2.944 | 1 | Glu101 | ~2.046 |
| **11.** | PelB | Taraxerone | -2.724 | Nil | Nil | Nil |
| **12.** | PelB | Taraxerol | -3.756 | Nil | Nil | Nil |

### 3.5. Trajectory analysis of MD simulations

A multifaceted computational tool entitled molecular dynamics (MD) determines and evaluates atomic particles’ physical movements in the context of macromolecular structure and function links. For a particular amount of time, the atoms and molecules were granted the ability to interact, reflecting how complicated the evolution of the system was [22]. The robustness of the assembled complex with compound and receptor structural rearrangements has been assessed by implementing a 100 ns MD simulation. The constancy and fluctuations of the Holo and Apo systems (Holo1: the RelA-Taraxerol complex, Holo2: the RelA-Taraxerol complex, Holo3: Taraxerone complex - SpoT, Holo4: Taraxerol complex - SpoT Holo5: Taraxerone complex of PqsA, Holo6: Taraxerol complex of PqsA, PqsR-Taraxerol complex in Holo8 and PqsR--Taraxerone complex in Holo7. To comprehend the fluctuating behavior and mode of binding, the subsequent compounds (Holo9: PelA-Taraxerone complex, Holo10: PelA-Taraxerol complex, Holo11: PelB-Taraxerone complex, and Holo12: PelB-Taraxerol complex) were evaluated using the Desmond suit (Schrödinger Release 2022-4: Maestro, Schrödinger, LLC, New York, NY, 2022). The dynamic resilience of both systems (Apo and Holo) was ascertained using the RMSD profile of the backbone atoms at 100 ns (Fig. 2A-2L). After 60 ns of MD simulations, the backbone RMSD graph of most of the Holo states confirmed an uninterrupted progression when juxtaposed with the Apo state.

**Figure. 2:**
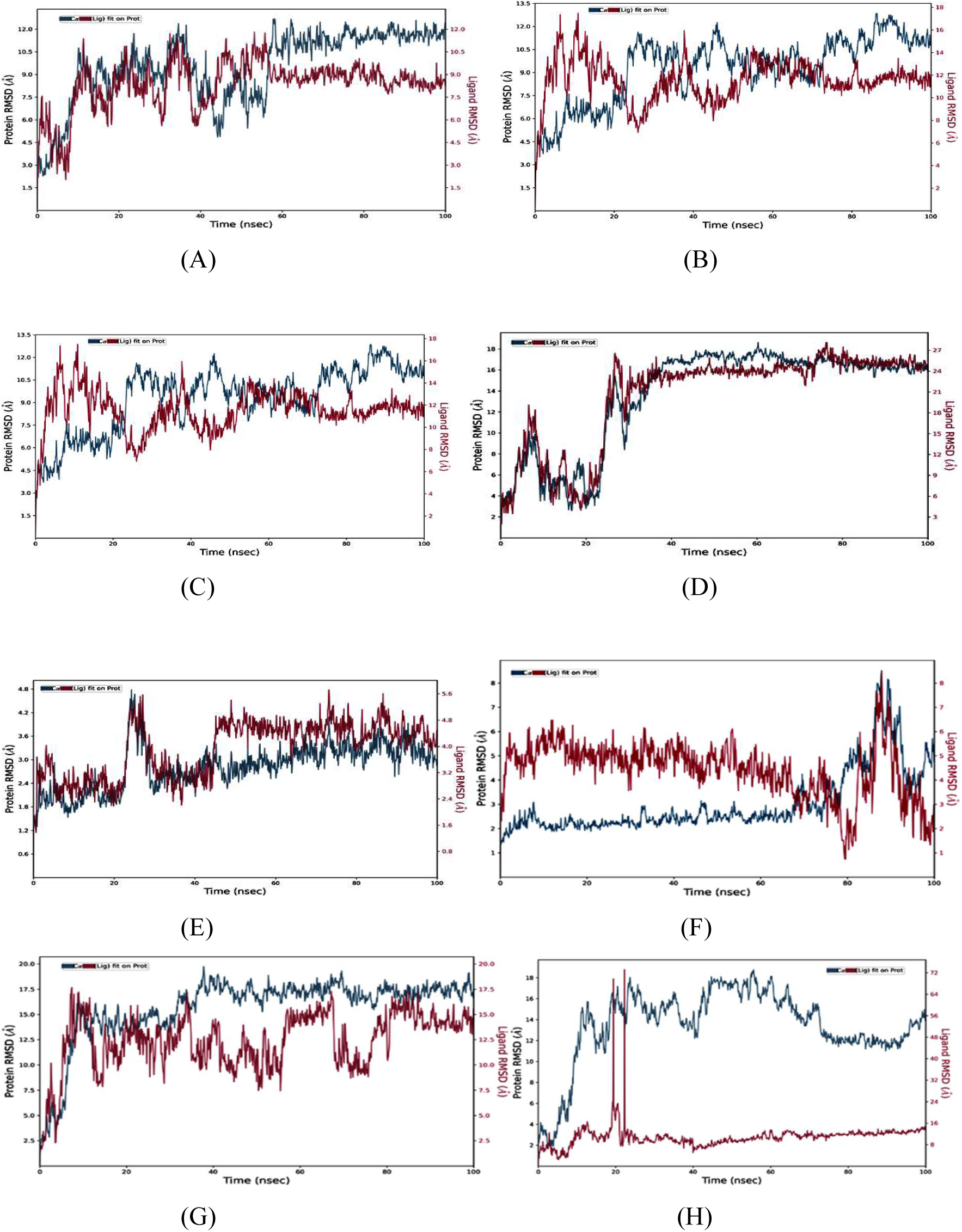
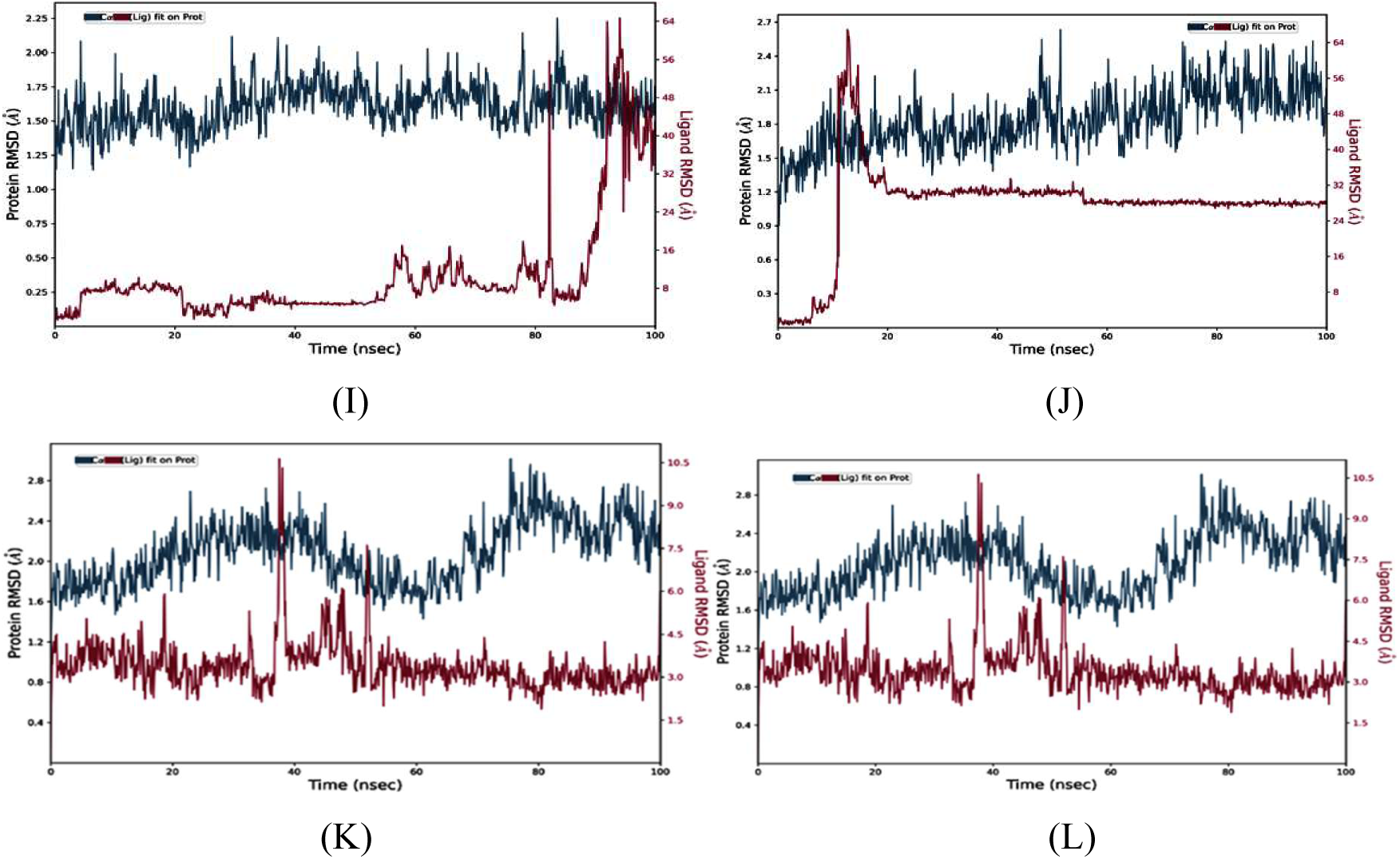
Conformational stability of Apo and Holo states of various proteins throughout 100 nanoseconds (ns) time period of MDS (**A)** Backbone-RMSD of RelA-Taraxerone complex **(B)** Backbone-RMSD of RelA - Taraxerol complex (**C**) Backbone-RMSD of SpoT - Taraxerone complex **(D)** Backbone-RMSD of SpoT - Taraxerol complex, **(E)** Backbone-RMSD of PqsA - Taraxerone complex, **(F)** Backbone-RMSD of PqsA – Taraxerol complex, **(G)** Backbone-RMSD of PqSR- Taraxerone complex, (H) Backbone-RMSD of PqSR-Taraxerol complex, (I) Backbone-RMSD of PelA- Taraxerone complex, (J) Backbone-RMSD of PelA -Taraxerol complex, (K) Backbone-RMSD of PelB- Taraxerone complex, (L) Backbone-RMSD of PelB-Taraxerol complex.

Compared to its Apo state, the backbone RMSD graph of the Holo1 state: RelA-Taraxerol complex confirmed a stable itinerary after 60 ns of simulation time. The Holo2 state: RelA-Taraxerol complex revealed a steady trajectory after 80 ns. In the MD simulations, the Holo1 state implied an RMSD value that varied from ∼7.6 to ∼10.1 Å for 60 to 100 ns, while the Holo2 state exemplar demonstrated an RMSD value extending from ∼7.8 to ∼10.0 Å for 80 to 100 MD simulation instances. This indicates that in preference to Taraxerone, the chemical Taraxerol may inhibit RelA and assist in stabilizing by disrupting its conformation. The RMSD graphs of Holo3: SpoT- Taraxerone complex, depicted inconsistent deviations during MD simulations, whereas Holo4: SpoT-Taraxerol complex depicted a stable trajectory after 80ns of time frame. Holo3 state reflected RMSD value between ∼2.9 to ∼6.0 Å Å from 90 to 100 ns of MD simulations whereas Holo4 state depicted a value between ∼15.9 to ∼17.0 Å from 80 to 100 of MD simulations time period. In case of Holo5: PqsA-Taraxerone complex, a stable trajectory was observed after 60 ns with RMSD values value between ∼3.3 to ∼4.7 Å whereas in Holo6: PqsA-Taraxerol complex an inconsistent deviations are seen with RMSD values ranging from ∼3.5 to ∼6.0 Å during 30 to 60 ns of MDS time. Holo7: PqsR-Taraxerone complex, depicted a stable trajectory after 85ns of MDS time frame whereas Holo8: PqsR-Taraxerol complex depicted a stable trajectory after 60ns with RMSD values, between ∼12.5 to ∼15.5Å and ∼3.8 to ∼4.0 Å respectively. Holo9: PelA-Taraxerone complex depicted a stable trajectory from 20 to 55ns of simulation time frame with RMSD values ∼0.20 to ∼0.25 Å, but after 55 ns inconsistent deviations were observed. In case of Holo10: PelA -Taraxerol complex, a steady trajectory after 60ns of MDS time frame with RMSD value ∼0.98 to ∼1.00 Å. Holo11: PelB- Taraxerone complex, and Holo12: PelB-Taraxerol complex depicted a stable trajectory after 60ns of MDS time period with RMSD values between ∼0.6 to ∼1.2 Å (Holo11), and ∼0.6 to ∼1.2 Å(Holo12) respectively. Taking together the RMSD values of all the Holo states it could be depicted that the compound Taraxerol could more potentially inhibit the targeted proteins compared to Taraxerone and can assist in stabilizing through changing its conformation.

Root mean square fluctuation (RMSF), which monitors residue swings, provided extra evidence for the RMSD result. Applying RMSF plots, the mobility among different residues was apparent in every circumstance (Fig. 3A-3L).

**Figure. 3.**
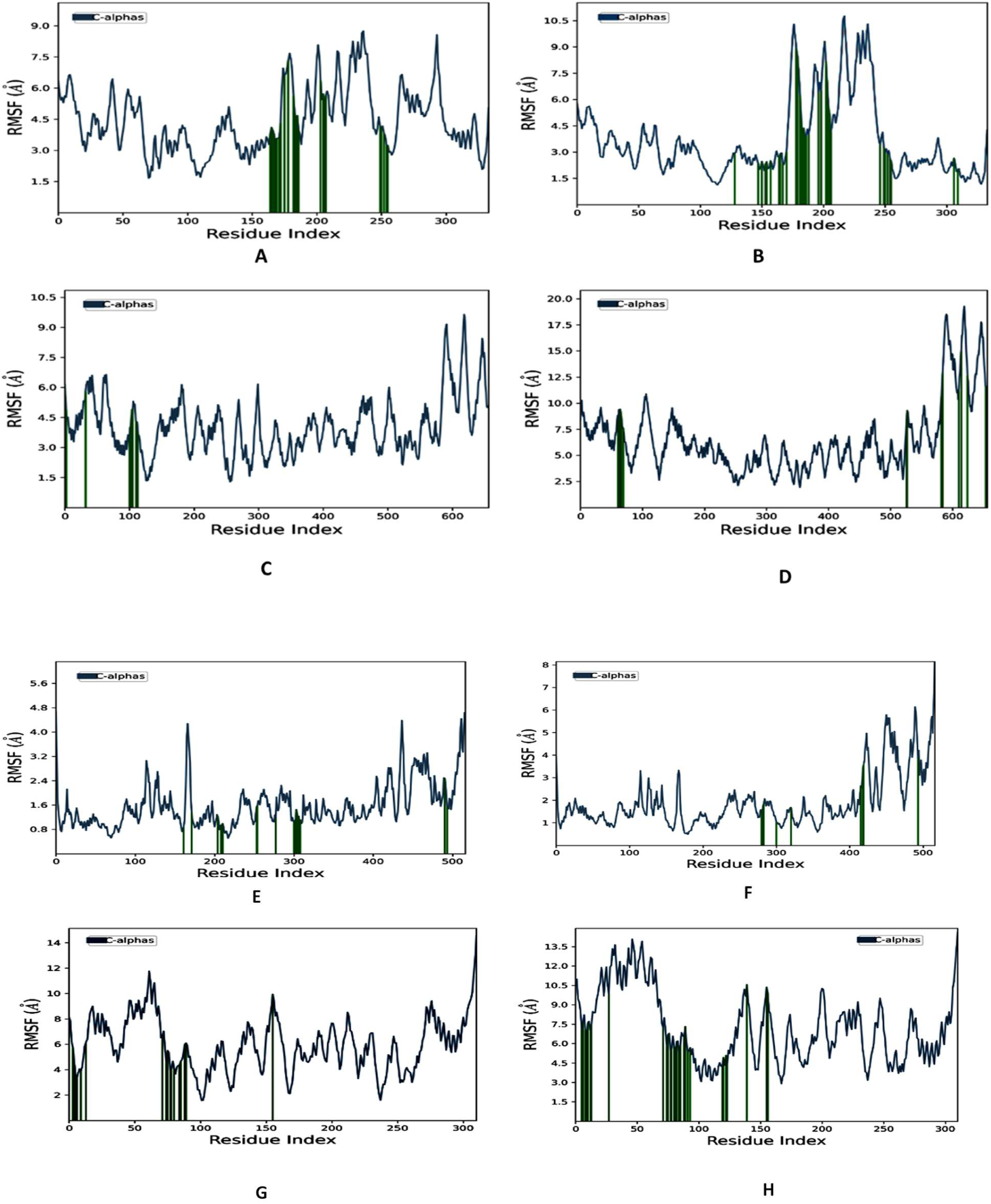
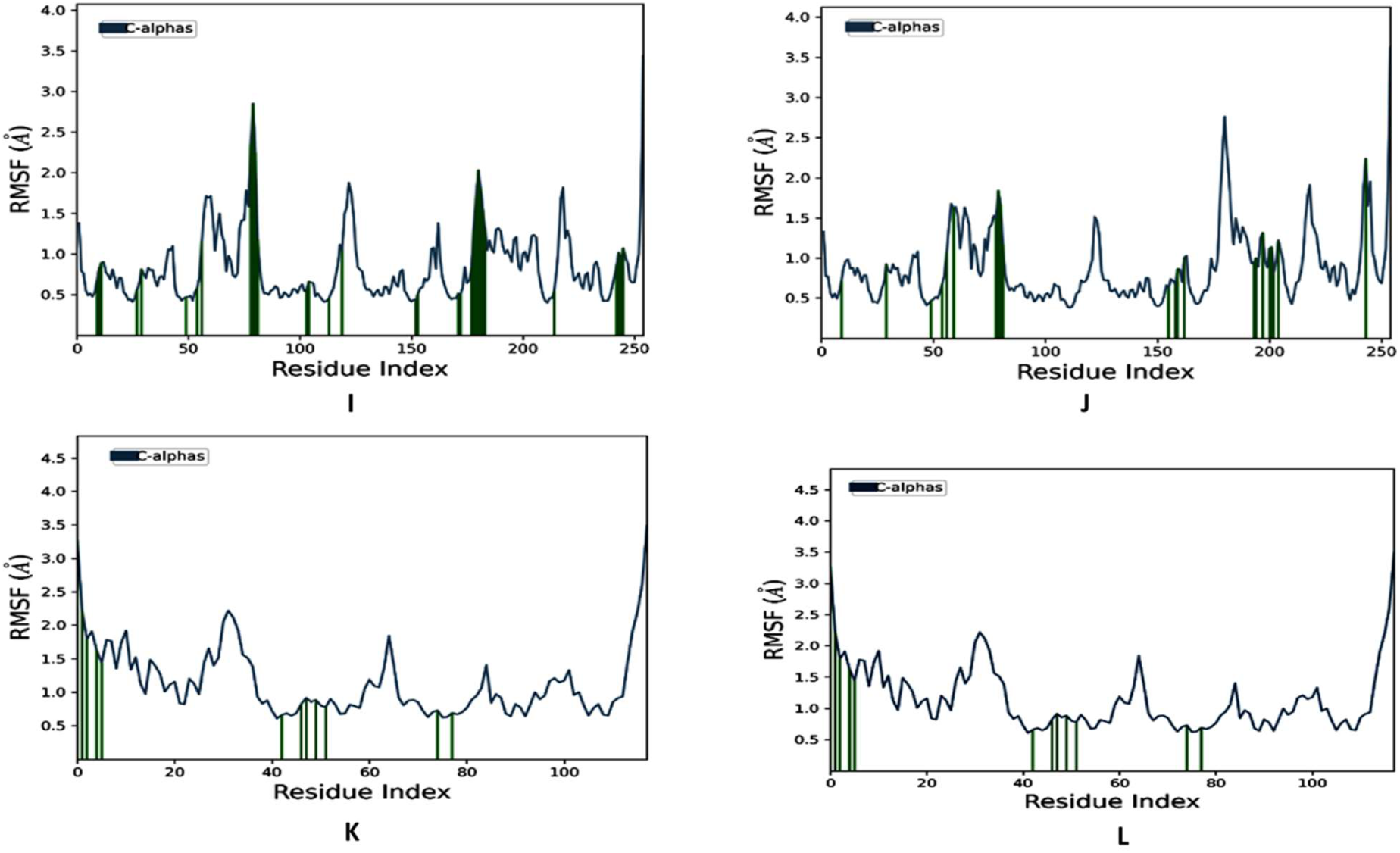
Conformational stability of Apo and Holo states of **various proteins** throughout 100 nanoseconds (ns) time period of MDS. Cα-RMSF profile of (**A**) RelA-Taraxerone complex**(B)** RelA - Taraxerol complex (**C**) SpoT - Taraxerone complex (D) SpoT - Taraxerol complex, (E) PqsA - Taraxerone complex, (F) PqsA – Taraxerol complex, (G) PqsR-Taraxerone complex, (H) PqsR- -Taraxerol complex, (I) PelA- Taraxerone complex, (J) PelA-Taraxerol complex, (K) PelB- Taraxerone complex, (L) PelB-Taraxerol complex.

The interaction between ingredients Taraxerone and Taraxerol might have prompted the amino acid residues in most of the Holo states to demonstrate more significant distinctions in their Cα atoms than other sites. The amino acid residues between 230 to 240 (Holo1,Holo2) and 580- 610 (Holo3,Holo4) ; 460-470 in Holo5,Holo6 ; 30-60(Holo7,Holo8); 120-125 (Holo9), 180-190 in Holo10; 28- 32 (Holo11, Holo12), displayed larger variations in their Cα atoms compared to other locations. In all states, approximately 10 terminal residues from the C- and N-terminal ends exhibited more extensive modifications, which can be eliminated. Green vertical bars signify protein residues that have an association with the ligand. This plot suggests that the mobility of residues in the Holo state may be shorter than that of the Apo state due to drug or ligand binding.

Attributes such as radius of gyration (yr) have been utilized to quantify the overall compactness for the stability of Taraxerone and Taraxerol in the binding pocket regardless of the receptor over the 100 ns simulation, as depicted in Fig. 4A-4L. A ligand’s primary moment of inertia, or radius of gyration (rGyr), measures its “extendedness,” whereas the number of internal H-bonds within a ligand molecule is conveyed in rGyr.

**Figure. 4:**
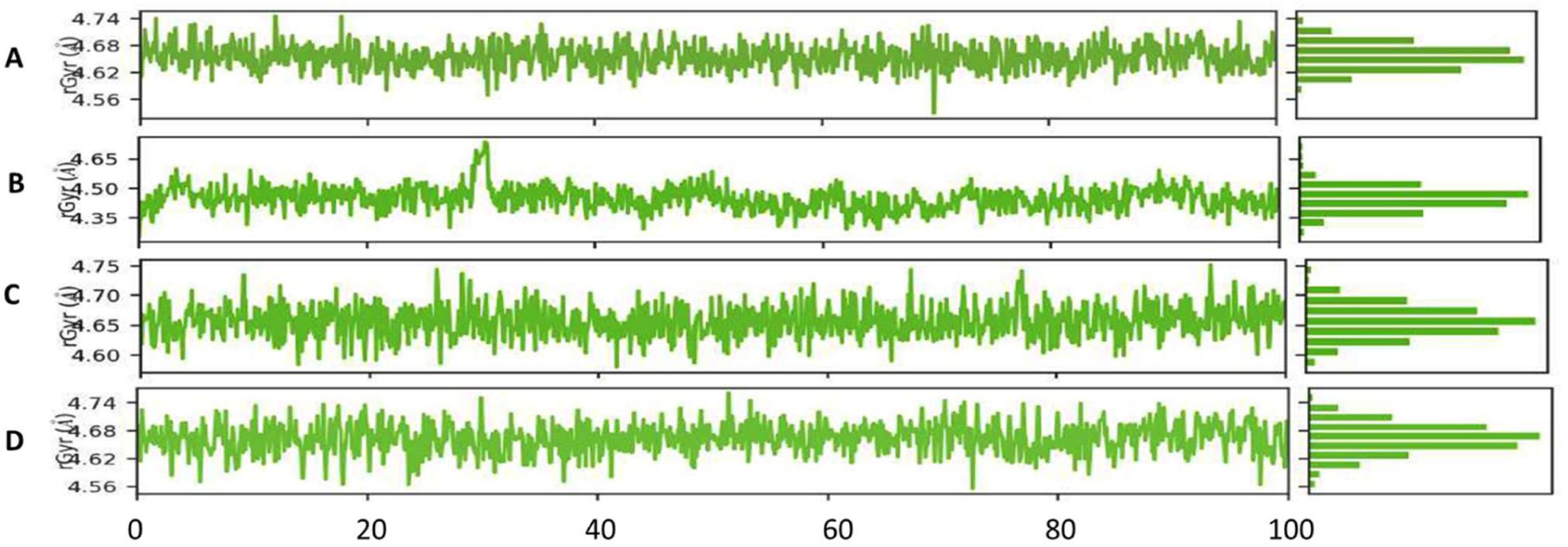
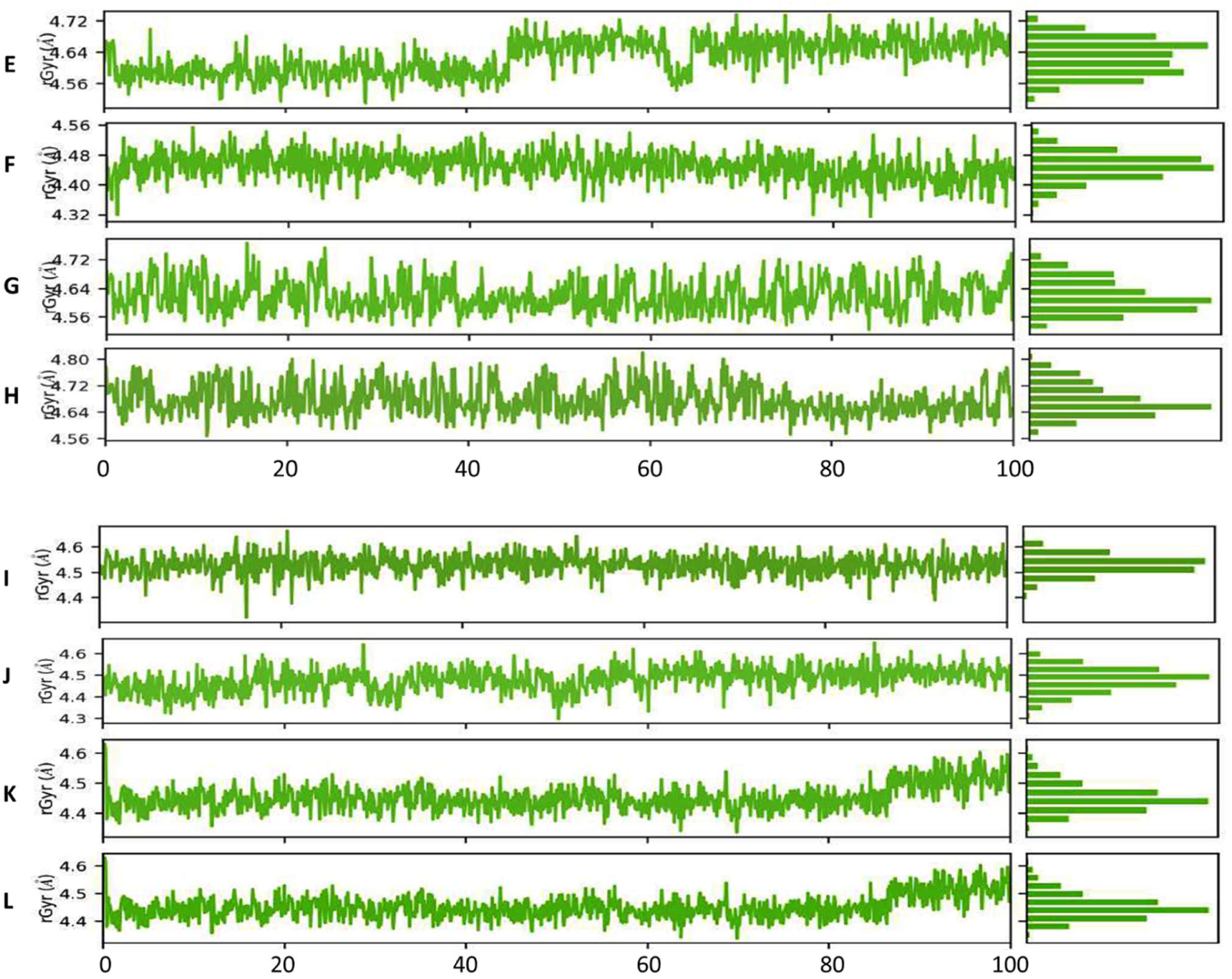
Conformational stability of **various proteins** throughout 100 nanoseconds (ns) time period of MD simulations. Radius of gyration (Rg) profile of (**A**) RelA-Taraxerone complex**(B)** RelA - Taraxerol complex (**C**) SpoT - Taraxerone complex (D) SpoT - Taraxerol complex, (E) PqsA - Taraxerone complex, (F) PqsA – Taraxerol complex, (G) PqsR-Taraxerone complex, (H) PqsR- -Taraxerol complex, (I) PelA- Taraxerone complex, (J) PelA -Taraxerol complex, (K) PelB- Taraxerone complex, (L) PelB-Taraxerol complex.

In each instance of Holo1 through Holo4, the fluctuation graphs of rGyr vs. simulation duration indicate rGyr stays constant, all while the simulation navigates after 80 ns. Around the receptor binding pocket of the protein RelA, the compound’s rGyr variation varied between ∼4.62 to ∼4.74Å in Holo1, confirming consistent Taraxerone functionality in all 80ns to 100ns of MD simulation. On the contrary, Holo2 reflected rGyr values ∼4.35 to ∼4.50Å, lower than Holo1. Compactness inversely pertains to rGyr value and vice versa, with lower rGyr values illustrating more accurate compactness [20], [21]. In the case of Holo3 and Holo4 the rGyr depicted values ∼4.60 to ∼4.75Å and ∼4.56 to ∼4.68 Å respectively. Here again the protein complexed with Taraxerol reflected a stable behaviour with more compactness compared to Holo3. Similarly, Holo5 to Holo8 represented a stable rGyr graph after 80ns of simulation period. The rGyr values are ∼4.60 to ∼4.72 Å for Holo5, ∼4.32 to ∼4.48 Å (Holo6), ∼4.56 to ∼4.72 Å (Holo7) and ∼4.57 to ∼4.80 Å (Holo8). Holo5 represented the lower rGyr values compared to Holo6, Holo7 and Holo8.

Subsequently, in case of Holo9 to Holo12, the rGyr values were ∼4.4 to ∼4.6Å for Holo9, ∼4.4 to ∼4.5 Å (Holo10), ∼4.4 to ∼4.6Å (Holo11) and ∼4.42 to ∼4.6 Å (Holo12) from 80ns to 100ns of MDS time frame. The protein complexed with Taraxerol reflected a stable behaviour with more compactness compared to Taraxerone. The analysis of rGyr values were well corroborated by RMSF analysis provides strong support for these rGyr results.

Their hydrophobic interactions regulate the way one specific solvents exhibit amino acids. The accessible surface area correlates precisely with the frequency of these interactions with the solvent and core protein residues. A decrease in the available solvent surface in the Holo states is demonstrated in the Solvent Accessible Surface Area (SASA) sketch (Fig. 5A-5L). Due to the amino acid residue migrating from the accessible to the buried region, the chemicals Taraxerone and Taraxerol can attach to proteins and change the hydrophilic and hydrophobic interface regions. This could have an impact on the alignments of the protein surface. During 80ns to 100ns MD simulation, the SASA graphs most of the Holo states reflected a buried state. The Holo1 state represented SASA with ∼140 to ∼245 Å^2^, whereas in the case of Holo2 the SASA value was ∼10 Å^2^ to ∼150Å^2^. Similarly, in Holo3 & Holo4, represented SASA with ∼290 Å^2^ to ∼310Å^2^ & ∼10 Å^2^ to ∼240Å^2^. This indicates that all across the 60–100 ns MD simulation period, the amino acid residues of Holo1 and Holo3 may move from the accessible to the buried region, triggering an orientational change in the protein surface. In case of Holo5 to Holo8, the SASA values were ∼80 Å^2^ to ∼150Å^2^(Holo5), ∼ 220 Å^2^ to ∼400Å^2^ (Holo6 ), ∼240 Å^2^ to ∼370Å^2^ (Holo7) and ∼160 Å^2^ to ∼290Å^2^ (Holo8) which depicted Holo8 with least SASA values reflecting its preference towards buried region compared to other Holo states. Subsequently, Holo9 to Holo12 the SASA values were ∼280 Å^2^ to ∼610Å^2^(Holo9), ∼ 280 Å^2^ to ∼290Å^2^ (Holo10), ∼10 Å^2^ to ∼300Å^2^ (Holo11) and ∼10 Å^2^ to ∼250Å^2^ (Holo12). Compared with the other Holo states, Holo10 was initially found upon moving from the approachable to the buried coverage zone.

**Figure. 5:**
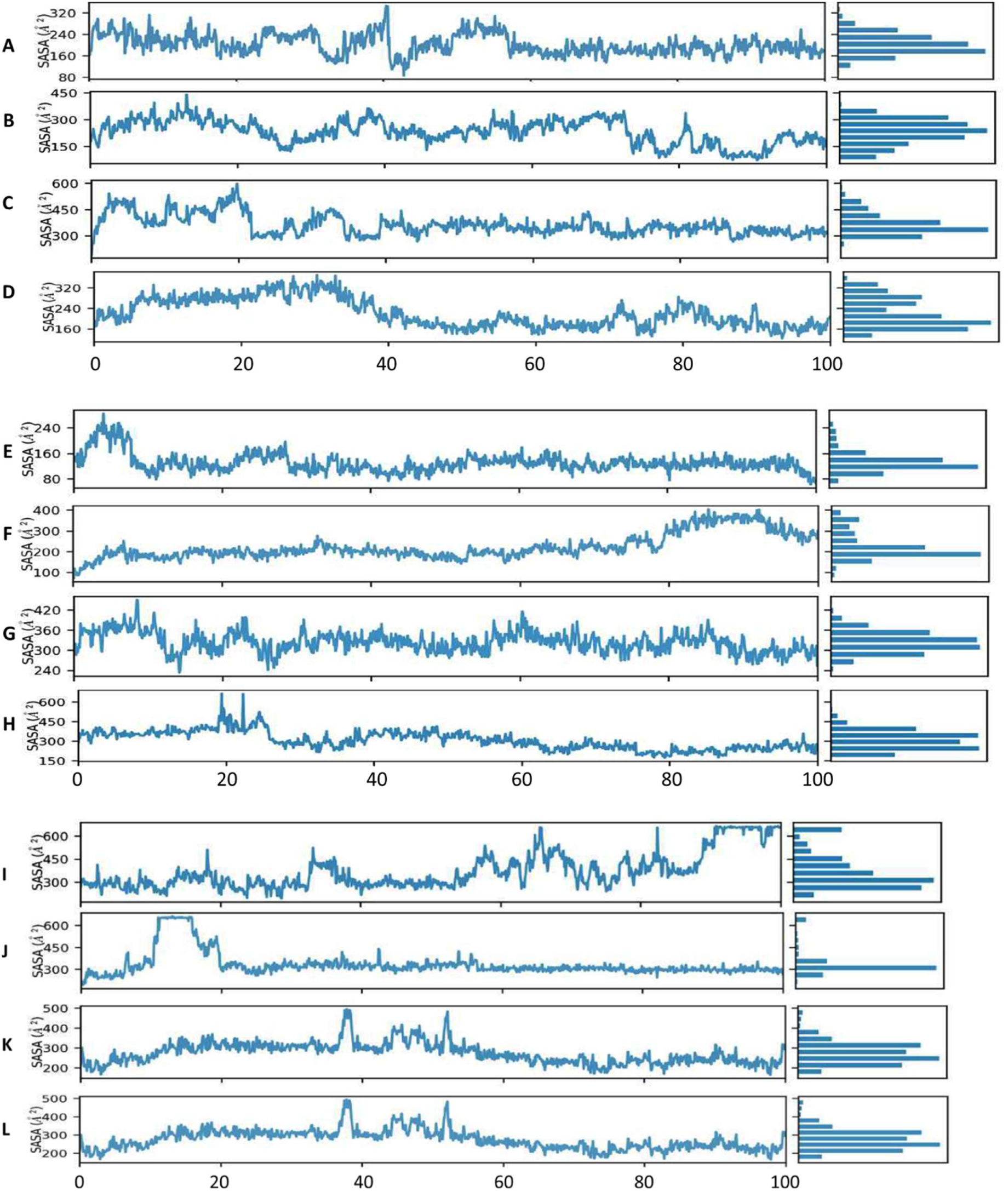
Solvent accessible surface area (SASA) analysis of **various proteins** throughout 100 nanoseconds (ns) time period of MD simulations. ((**A**) RelA-Taraxerone complex**(B)** RelA - Taraxerol complex (**C**) SpoT - Taraxerone complex (D) SpoT - Taraxerol complex, (E) PqsA - Taraxerone complex, (F) PqsA – Taraxerol complex, (G) PqsR- Taraxerone complex, (H) PqsR- -Taraxerol complex, (I) PelA- Taraxerone complex, (J) PelA -Taraxerol complex, (K) PelB- Taraxerone complex, (L) PelB-Taraxerol complex.

### 3.6. H-bond analysis

Schrödinger Release 2022-4 was used to illustrate the intermolecular hydrogen bonds of the Holo states during MD simulations (supplementary Fig. S1 A-1AJ). Components of intermolecular hydrogen bonds were discovered during the simulation of all Holo states. For Holo1 and Holo2, the post-MD simulation scrutiny indicated that there was indeed no H-bond. Throughout the simulation period, the stability of the drug-target complex corresponded precisely with the total quantity of H-bonds. The Holo1 state’s intermolecular hydrogen bonds were monitored (Fig. S1 A). The stacked bar chart of Holo1 in Fig. 2A indicates that leu582, one of the amino acid residues of RelA, may be crucial for the binding and inhibition of the protein. These are the most critical amino acid residues for protein binding and activity. Specific protein residues could participate across multiple interactions of the same subtype with the ligand; this histogram can demonstrate values more than 0.05. Throughout the simulation, the overall number of intermolecular hydrogen bonds was represented in contradiction by the Holo1 state simulation (Fig. S1B). In the scenario of the post-MD of Holo1; no H-bond was shown (Fig. S1C). The stability of the drug-target complex was directly correlated with the number of H-bonds during the simulation. H-bond forming residues like Gly598 were broken down in Holo1 simulations; however, these connections were later compensated for by fresh hydrophobic contacts and van der Waals interactions (Fig. S1C). Holo2’s stacked bar chart in Fig. S1D confirms that amino acid residues of RelA, including Gln571 and Asp578, may be essential for the binding and control of the protein. Certain protein residues might participate across multiple interactions of the same subtype with the ligand; this histogram may show values in excess of 0.05. Throughout the simulation, the total number of intermolecular hydrogen bonds was regularly reflected by the Holo2 state simulation (Fig. S1E). When Holo2 was post-MD, no H-bond was represented. A few new hydrophobic interactions and van der Waals interactions were used to make up for the breakage of the H-bond forming residue Gln581 amid the simulations of Holo2 (Fig. S1F).

Analogously, in Holo3 and Holo4, the stacked bar chart of Holo3 in Fig. S1G indicates that amino acid residues like Tyr45 and Val149 might be significant in protein binding and regulation. Values reaching 0.10 in this histogram are possible due to the possibility of multiple interactions of the same subtype entre some protein residues and the ligand. Throughout the simulation, the total number of intermolecular hydrogen bonds was represented sporadically by the Holo3 state simulation (Fig. S1H). When Holo3 was post-MD, no H-bond was represented. H-bond forming residues like Thr47 were broken down in initial Holo3 simulations. Still, later on, van der Waals interactions and novel hydrophobic interactions made up for it (Fig. S1I). In the context of Holo4, the amino acid residues Phe108 and Lys111 may be essential for the binding and regulation of the protein, demonstrating by the stacked bar chart in Fig. S1J. Values above 0.1 in this histogram are possible due to various interactions of the same subtype between some protein residues and the ligand. Throughout the simulation, the quantity of intermolecular hydrogen bonds was consistently reflected by the Holo4 state simulation (Fig. S1K). When Holo4 was post-MD, no H-bond was represented. H-bond forming residues like Met106 were disrupted during Holo4 simulations, but consequently, van der Waals interactions and new hydrophobic interactions formed the basis for it (Fig. S1L).

The amino acid residues Gln162 and Tyr211 are represented in the stacked bar chart of Holo5 in suppl. Fig. 1M and they may be significant in the binding and control of the protein. Throughout the simulation, there were multiple instances of intermolecular hydrogen bonding (Fig. S1N). In the specific case of the post-MD of Holo5 (TYR211), one H-bond was displayed. H-bond-delivering residues like TYR304 were broken down through Holo5 simulations, but afterward, new H bonds, hydrophobic acquaintances, and van der Waals interactions were generated (suppl. Fig.1O). Addressing Holo6, the stacked bar chart (suppl. Fig. 1P) indicated that Pro281, an amino acid residue, would be crucial in the binding and regulation of the protein. Throughout the simulation, there were additionally varying numbers of intermolecular hydrogen bonding (suppl. Fig. 1Q). In this particular instance of the post-MD of Holo6, one H-bond was represented (PRO281). Despite H-bond offering residues like PRO281 failed to break down in Holo6 simulations, they were later compensated by novel hydrophobic relationships and van der Waals interactions (Fig. S1R). The H-bond forming residue (PRO281) in Holo6 (PqsA – Taraxerol complex) failed to compensate for the few new interactions. This indicates that Taraxerol is effective against the targeted protein PqsA since PRO281 may be a necessary residue that prevents Quorum sensing from activating.

The amino acid residues Leu6, Leu75, and Asn89 in Holo7 were apparent in the stacked bar chart (Fig. S1S), implying that these residues may be significant in protein binding and regulation. Throughout the simulation, there were numerous varying numbers of intermolecular hydrogen bonding (Fig. S1T). When Holo7 was post-MD, no H-bond was depicted after the disintegration of H-bond-forming residues like ASN76, new H bonds, hydrophobic interactions, and van der Waals interactions were established up for (suppl.Fig. 1U). Asn7, Gln78, Gln79, Asn89, and Asn 120 amino acid residues are highlighted in the stacked bar chart (suppl. Fig. 1V) for Holo8, which indicates that the residues in question may be significant in protein binding and regulation. A steady proportion of hydrogen bonds exists between molecules across the simulation (Fig. S1W). In the case of the post-MD of Holo8, two H-bonds have been identified (Asn89, Asn 120). In Holo8 simulations, Asn76 and other H-bond-forming residues suffered disruptions, but eventually, new H-bonds, hydrophobic interactions, and van der Waals acquaintances were made up for (Fig. S1X). The amino acid residues Trp126 and Trp224 have been illustrated in the stacked bar chart (Fig. S1Y) in Holo9 and may be substantial in protein binding and regulation. Throughout the simulation, there were additionally varying numbers of intermolecular bonds involving hydrogen (Fig. S1Z). When Holo9 was post-MD, no H-bond was depicted. H-bond forming residues like TRP224 experienced disruption during Holo9 simulations, but consequently, novel hydrophobic interactions and van der Waals interactions have been lined up for (Fig. S1 AA). The amino acid residues Glu101, Asp103, and Gly224 in Holo10 have been illustrated in the stacked bar chart (Fig. S1AB), which hints that these residues may be significant in protein binding and regulation. Throughout the simulation, there were numerous instances of intermolecular hydrogen bonding (Fig. S1AC). When Holo10 was post-MD, no H-bond was implied. H-bond forming residues, such as GLU101, suffered interference during Holo9 simulations; after that, fresh hydrophobic interactions and van der Waals contacts were established (Fig. S1AD).

Similarly, the stacked bar charts (Fig. S1AE) for Holo11 and Holo12 point out that amino acid residues like Glu319 and Asp320 (Holo11) may be essential to the binding and regulation of the protein. During the simulation, there is a fixed quantity of hydrogen bonds between molecules (Fig. S1AF). In Holo11’s post-MD instance, one H-bond appeared (Gln366). H-bond-forming residues encountered disruption during Holo11 simulations, but subsequently, novel H-bonds, hydrophobic interactions, and van der Waals interactions were set up (Fig. S1AG).

Alongside this, Holo12’s stacked bar chart (Fig. S1AH) reveals how amino acid residues like Glu319 and Asp320 could be critically important to the binding as well as regulation of the protein. A steady quantity of hydrogen bonds between molecules around the simulation (suppl.Fig.1AI). In the case of Holo12’s post-MD, one H-bond was represented (ASP392). H-bond-forming residues in Holo12 simulations were impacted, but alternative H-bonds, hydrophobic interactions, and van der Waals interactions emerged to make up for (Fig. S1 AJ).

## 4. Discussion

Absurd use of antibiotics triggers AMR in both clinical and environmental bacterial isolates. Strategies to combat drug resistance are majorly focused on inhibition mechanisms responsible for AMR spread. Often these strategies ignore the biofilm aspect of AMR in bacterial pathogens. For testing drug or phytochemical for its prospective use as an antibacterial, the target is always on growth inhibition, instead of biofilm inhibition. Alternate strategies to combat drug resistance can be accomplished by targeting biofilms. The findings of the present study support targeting multiple genes responsible for biofilm-related drug resistance should be considered. The virtual screening outcome of the present study revealed that all investigated compounds from mangrove plants exhibited better binding energies (Table S1) during its interaction with all 18 protein targets. Out of 18, the targets PqsA and PqsR, represented a better average with consistent interactions with most of the compounds. However, it binding with Taraxerone and Taraxerol displayed better binding affinity as compared to the other phytochemicals. These compounds have been reported for their antibiofilm and acyl homoserine lactone (AHL) quorum sensing inhibition activity, but PQS quenching activity has not been evaluated thoroughly [28]. Subsequently, the molecular docking depicted Taraxerol with the highest binding affinity against PqsA with -3.904 kcal/mol among all the six targets associated with persistence (RelA and SpoT), QS (pqsA, and pqSR), and EPS synthesis (PelA and PelB) in *P. aeruginosa*.

In *P. aeruginosa* biofilm development and resistance to antibiotic therapy is linked to multiple QS network. QS coupled biofilm regulation contributes to the pathogenicity of the bacterium leading to the treatment failure. In *P. aeruginosa* apart from iron scavenging, 2-heptyl-3-hydroxy-4(1H)-quinolone (PQS) regulates numerous virulence genes that create a global regulatory network responsible for the regulation of 10-12% of the function genome in the bacterium [29]. Interestingly PQS system works synergistically with other AHL-based QS systems in the bacterium and regulates virulence gene expression. PQS serves as a stress warning system shaping *P. aeruginosa* population structure in biofilm and its response to hostile environmental conditions e.g. antibiotic, pH, salinity, host immune system etc. [30], [31]. PQS could be a potential target to control virulence traits and host-microbe interactions [32]. In *P. aeruginosa* PqsR, a Lys-R type transcriptional regulator, with its cognate signal molecules 4-hydroxy-2-heptylquinoline (HHQ) and 2-heptyl-3-hydroxy-4-quinolone (PQS) controls the overall genome within the bacterium.

PqsR, is a multiple virulence factor regulator (MvfR), which also controls the formation of antibiotic-induced persister cells in *P. aeruginosa*. Thus, inhibition of the PqsR system can contribute to persistence inhibition [33]. Apart from PqsR system, the major regulators of persister cell formation are *relA* and *spoT. relA* mutant of *P. aeruginosa* has been reported with reduced virulence as compared to wild type. *relA* inhibition can increase phycocyanin production but negatively affect the elastase level. *P. aeruginosa* elastase enzyme has tissue-damaging activity important for localized infections. This enzyme is one of the best classified virulence-enhancing factors of the bacterium [34]. Attenuation of both *relA* and *spoT* leads to a 1000-fold reduction in cell survival upon antibiotic exposure, by downregulation of persister cells, elastase levels, and biofilm development [35], [36]. The results of docking are well aligned with the findings of molecular dynamic studies which indicated that both the compounds Taraxerone and Taraxerol displayed good binding affinity and stabilizing the complex against RelA and SpoT, indicating that these natural products could be potentially useful in controlling persister cell formation within biofilm and can be used as antibiotic potentiators.

Within the biofilm matrix, the cationic polymer designated as pellicle (Pel) polysaccharide simplifies communication between cells. Made from dimeric repetitions of galactosamine and N-acetylgalactosamine, it is an additionally de-N-acetylated linear polymer of α-1,4-N-acetylgalactosamine [37]. Pel polysaccharide is critical for virulence and persistence in *P. aeruginosa*. The pel operon consists of seven genes (*pelABCDEFG)* where *pelA* has glycosidic activity. *pelB* is a membrane-embedded porin, which is required for the deacetylation activity of *pelA*. Inhibition of *pelAB* activity is considered as a viable strategy to block biofilm development in *P. aeruginosa* [38], [39]. Identification of therapeutic molecules that can suppress virulence and AMR in biofilm by targeting biofilm matrix polysaccharides could be an innovative strategy to increase antibiotic efficiency by removing diffusion barriers within biofilm [40]. Therapeutic approaches targeting biofilm exopolysaccharides like Pel principally aim to disrupt EPS using compounds that act against exopolysaccharides. Strategies targeting EPS are designed to enhance the penetration of antibiotics into the biofilm [41]. In the present study, the computational investigations represented PelA and PelB as the potential targets for the compounds Taraxerone and Taraxerol based on their binding affinities that could inhibit the targets from EPS synthesis.

Additionally, the findings of this *in silico* study may serve as confirmation proof for further *in vitro* and *in vivo* investigations that are required to verify the anti-biofilm drug potency and efficacy of phytochemicals derived from mangrove plants, such as Taraxerone and Taraxerol, against proteins necessary for the synthesis of biofilm matrix, cell-cell communication, and the formation of persistence cells induced by antibacterial medications.

## 5. Conclusion

The application of phytochemicals as natural antimicrobial agents and antibiotic potentiators could be a promising strategy to control bacterial biofilm in both human and zoonotic infections. AMR in bacterial pathogens has aggravated the search for novel drugs derived from natural sources to control infection. However, its treatment becomes extremely challenging if there is an onset of biofilm formation. During the biofilm phase of growth, both genetic and biofilm microenvironments add toward antibiotic failure. Owing to the multiple therapeutic effects of phytochemicals, a single phytochemical can be harnessed to target all pathogens-related trends of biofilm.

*P. aeruginosa* is often studied as a model pathogen for biofilm studies because of its unique and tenacious nature. The bacterium has been routinely reported in a variety of infections and comes under WHO’s priority list of pathogens. In the present study, we aimed to evaluate natural compounds derived from *Avicenna* species for the inhibition of virulence-related genes in the bacterium. Our strategy aims to study three important traits of *P. aeruginosa* biofilm (i.e. PQS, persister cells and EPS), which are often ignored even though crucial for combatting AMR. A Plethora of *in silico* approaches has been studied earlier in search of novel antibiofilm natural compounds, but they only focus on AHL-based QS in the bacterium. Moreover, there are hardly any studies where EPS and persister cells are considered. All 49-chemical showed stable binging with 18 proteins that were targeted. The consensus results of virtual screening and docking studies were well collated with the MD simulations analysis and depicted the compounds Taraxerone and Taraxerol against six potential targets with QS (PqsA and PqsR), EPS synthesis (PelA and PelB) and persistence (RelA and SpoT) activity of *P. aeruginosa.* Inhibition of the above-mentioned targets by potential compounds like Taraxerone and Taraxerol could play a pivotal role in the arena of AMR. The present investigation could cut short many *in-vitro* and *in-vivo* experiments, hence it is important to acknowledge the facts of this *in silico* findings which is further need to be confirmed by *in vitro* and *in vivo* studies to evaluate their drug potency.

## Authors’ contributions

SB and NAK contributed to the designing of concepts, experiments, manuscript writing, data analysis, and editing. SB did all MD stimulations. YKM contributed to data analysis and manuscript writing.

## Ethics approval

Not Applicable

## Acknowledgment

We are thankful to the Gujarat Biotechnology University-Gandhinagar, National Institute of Pharmaceutical Education and Research-Ahmedabad, University of Science and Technology Meghalaya, and Chettinad Academy- of Research and Education-Kelambakkam for providing infrastructure and resources to carry out this work.

