## Supplementary material for "Multilevel computational approach to unlock the potential inhibitors of biofilm-EPS, persistence and quinolone signalling in *Pseudomonas aeruginosa* using mangrove-derived bioactive phytochemicals": suppl.

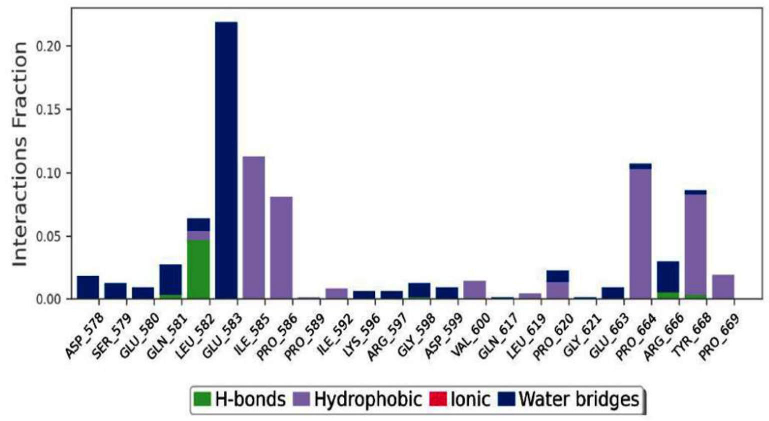


**(A)**

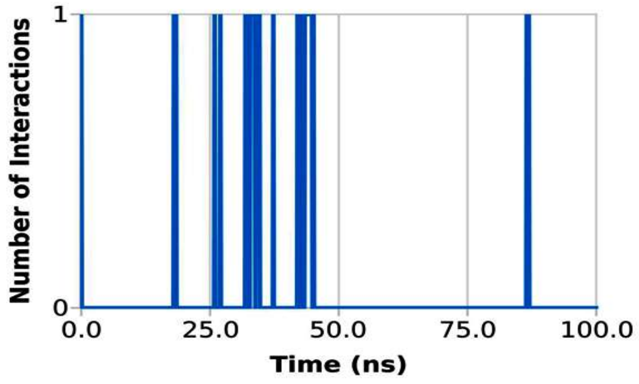


**(B)**

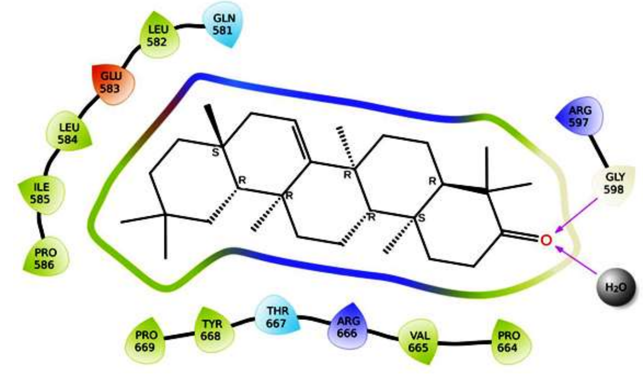


**(C)**

**Figure S1 (A-C):** Stacked bar chart showing protein ligand contacts plot for (A) RelA-Taraxerone complex during the simulation of 100 ns. (B) Deviation of H-bonds contributed in interaction during 100 ns simulation. (C) Post-MD simulations intermolecular hydrogen bonding, electrostatic and hydrophobic contacts formed between Post MDS RelA-Taraxerone complex**.** The image was generated by ligand interaction module of Schrödinger.

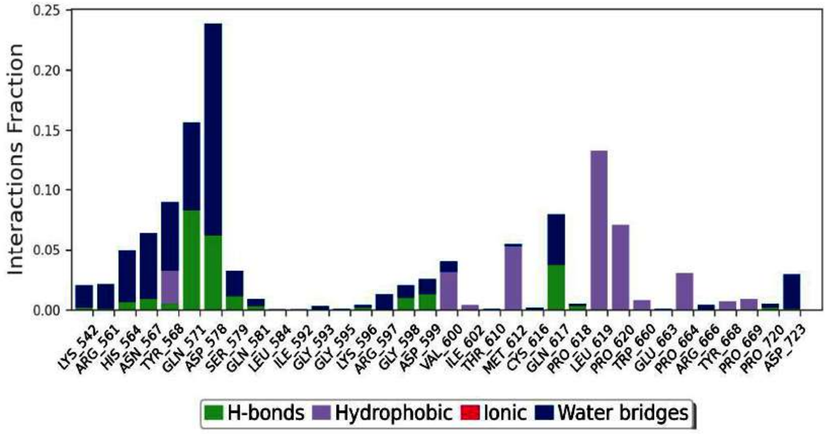


**(D)**

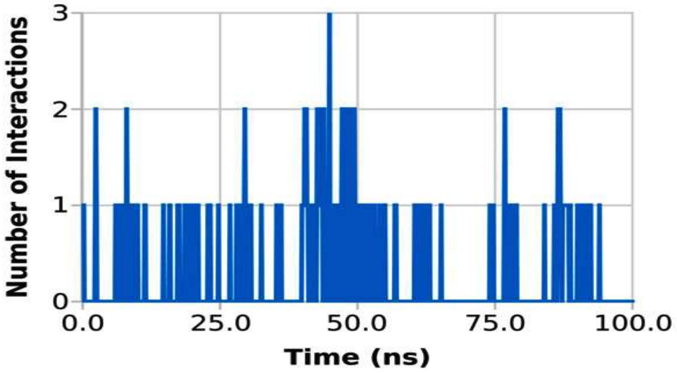


**(E)**

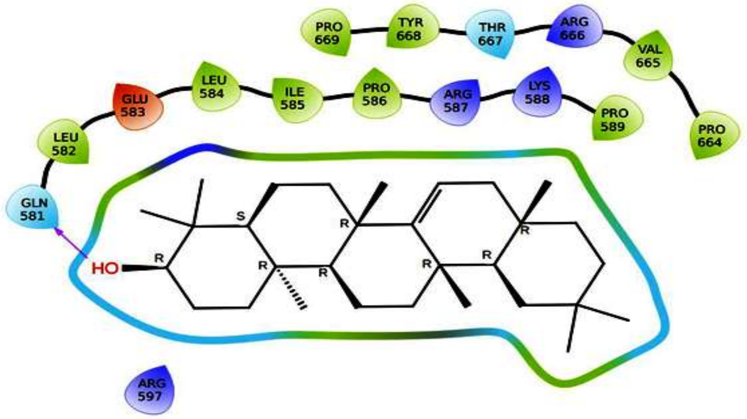


**(F)**

**Figure S1 (D-F)** Stacked bar chart showing protein ligand contacts plot for (D) RelA - Taraxerol complex during the simulation of 100 ns. (E) Deviation of H-bonds contributed in interaction during 100 ns simulation. (F) Post-MD simulations intermolecular hydrogen bonding, electrostatic and hydrophobic contacts formed between Post MDS RelA - Taraxerol complex**.** The image was generated by ligand interaction module of Schrödinger.

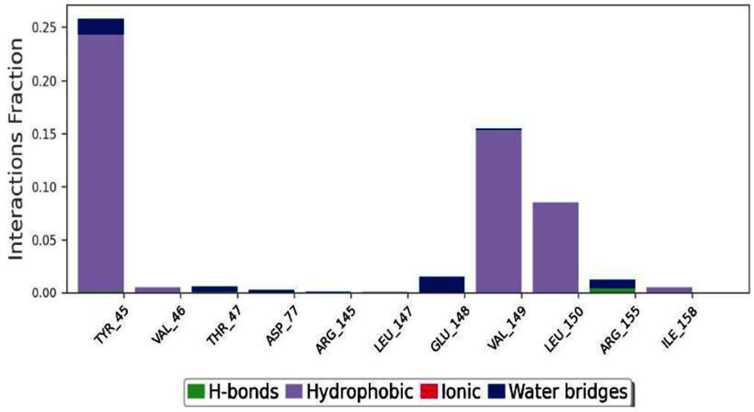


**(G)**

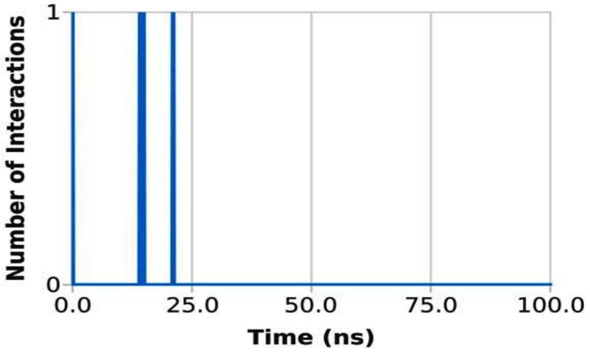


**(H)**

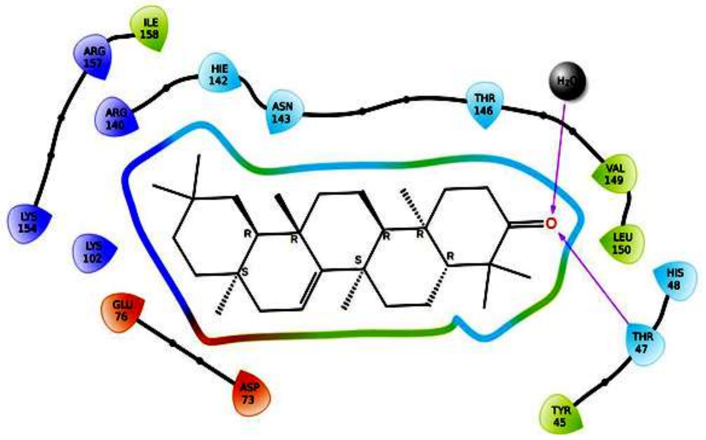

**(I)**

**Figure S (G-I):** Stacked bar chart showing protein ligand contacts plot for (G) SpoT - Taraxerone complex during the simulation of 100 ns. (H) Deviation of H-bonds contributed in interaction during 100 ns simulation. (I) Post-MD simulations intermolecular hydrogen bonding, electrostatic and hydrophobic contacts formed between Post MDS SpoT - Taraxerone complex. The image was generated by ligand interaction module of Schrödinger**.**

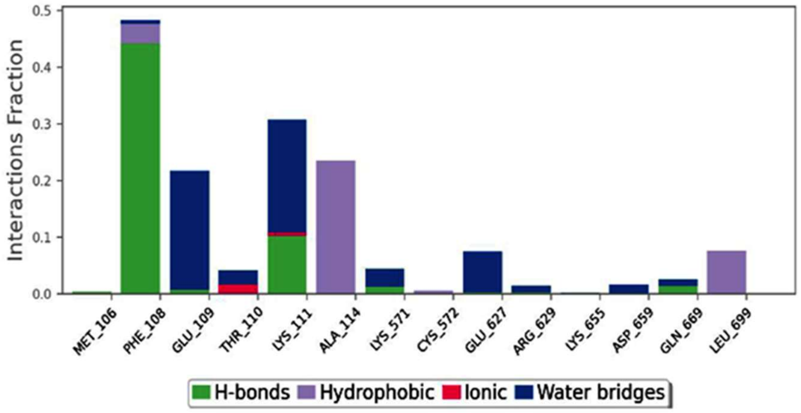


**(J)**

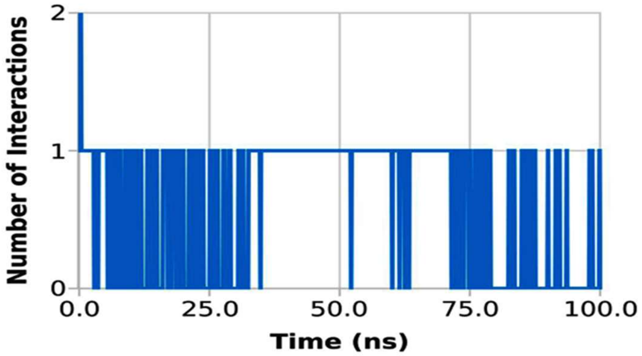


**(K)**

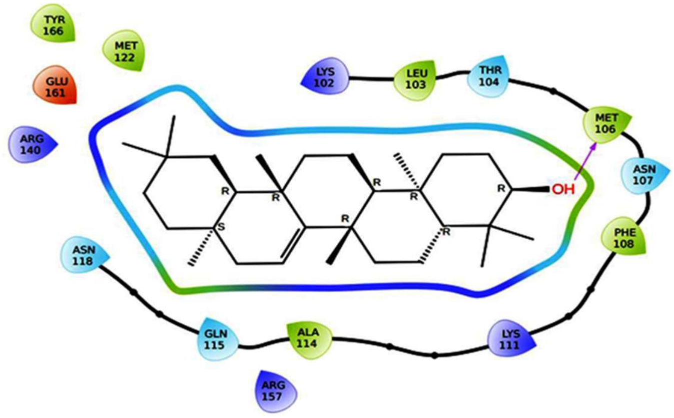


**(L)**

**Figure S1 (J-L):** Stacked bar chart showing protein-ligand contacts plot for (G) SpoT - Taraxerol complex during the simulation of 100 ns. (H) Deviation of H-bonds contributed in interaction during 100 ns simulation. (I) Post-MD simulations intermolecular hydrogen bonding, electrostatic and hydrophobic contacts formed between Post MDS SpoT – Taraxerol complex. The image was generated by ligand interaction module of Schrödinger**.**

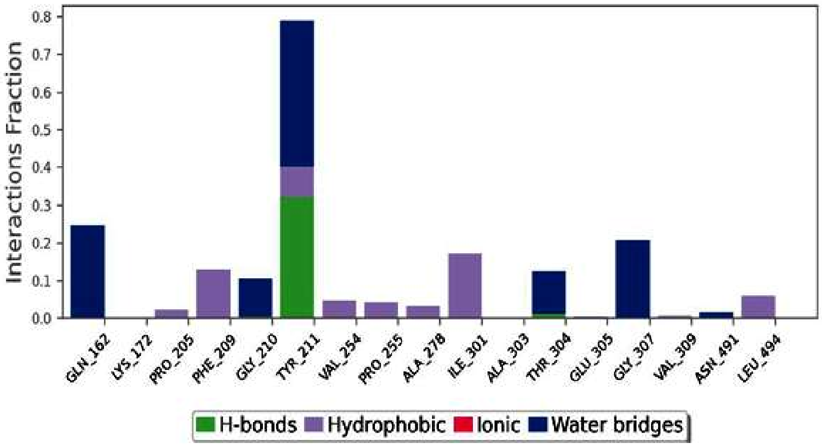


**(M)**

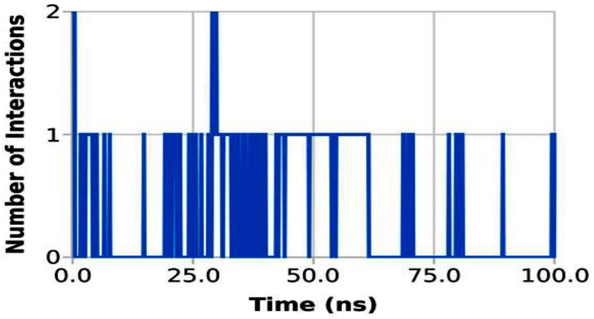


**(N)**

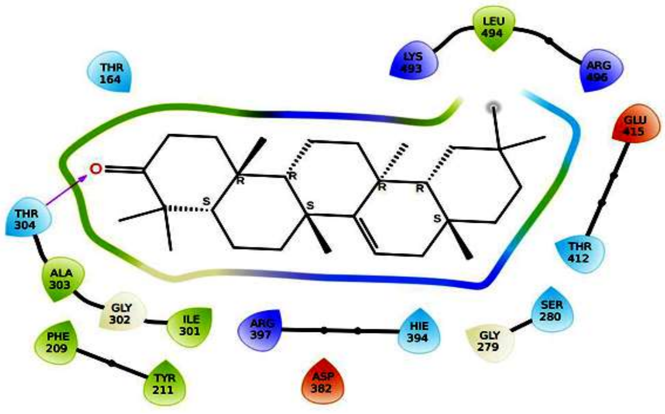


**(O)**

**Figure S1 (M-O):** Stacked bar chart showing protein ligand contacts plot for (M) PqsA - Taraxerone complex during the simulation of 100 ns. (N) Deviation of H-bonds contributed in interaction during 100 ns simulation. (O) Post-MD simulations intermolecular hydrogen bonding, electrostatic and hydrophobic contacts formed between Post MDS PqsA - Taraxerone complex. The image was generated by the ligand interaction module of Schrödinger.

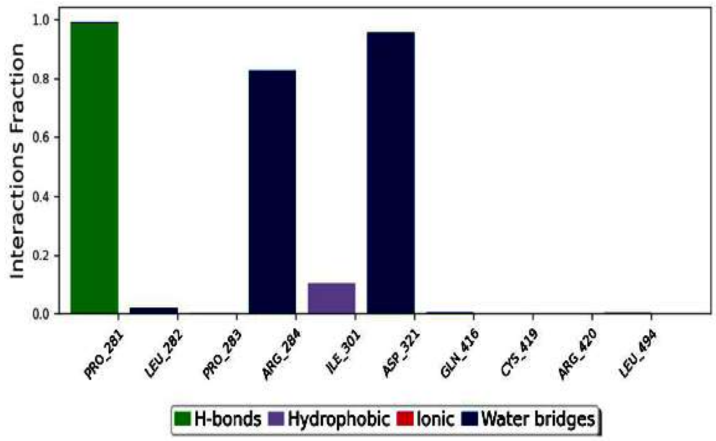


**(P)**

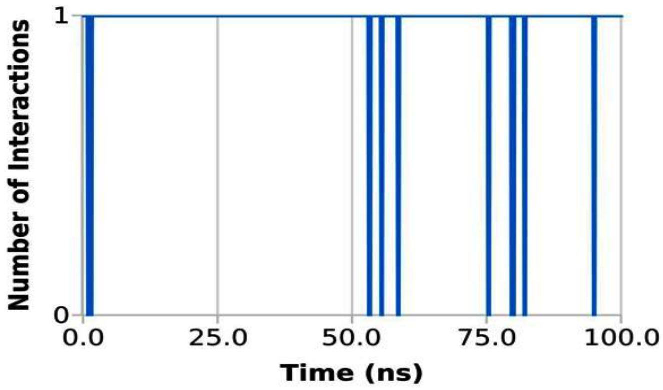


**(Q)**

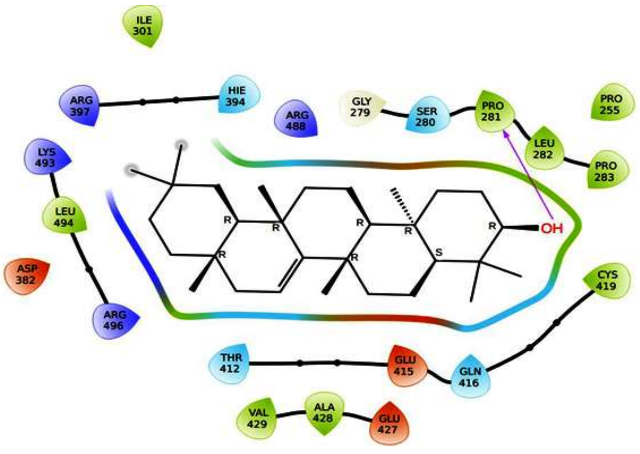

**(R)**

**Figure S1 (P-Q).** Stacked bar chart showing protein ligand contacts plot for (P) PqsA – Taraxerol complex during the simulation of 100 ns. (Q) Deviation of H-bonds contributed in interaction during 100 ns simulation. (R) Post-MD simulations intermolecular hydrogen bonding, electrostatic and hydrophobic contacts formed between Post MDS PqsA – Taraxerol complex**.** The image was generated by ligand interaction module of Schrödinger.

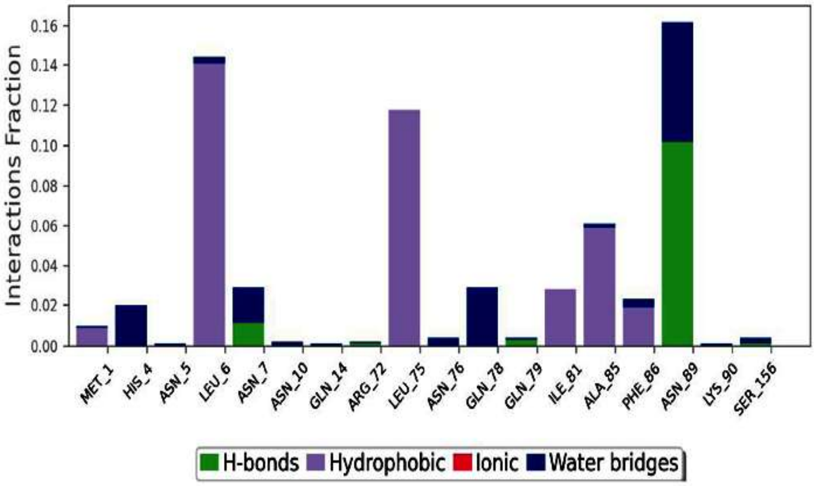


**(S)**

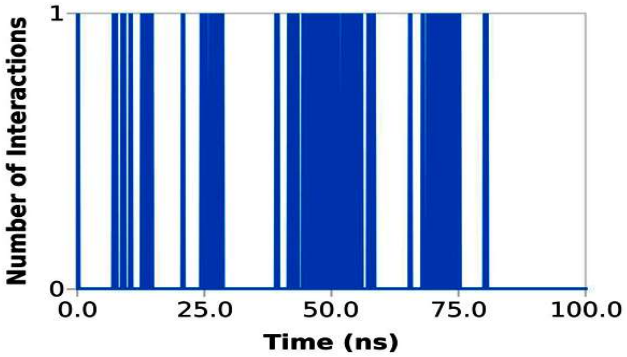


**(T)**

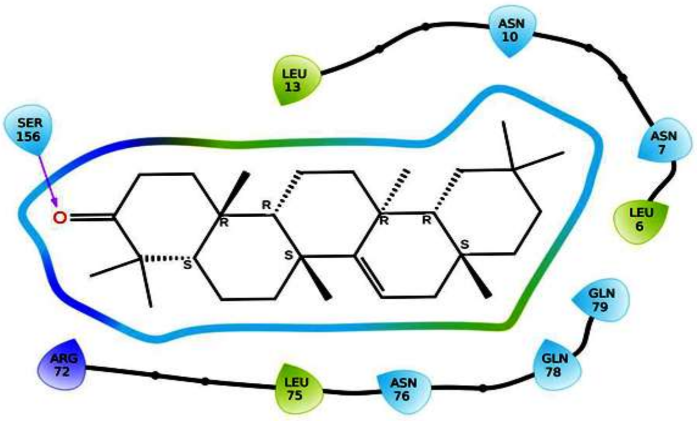

**(U)**

**Figure S1 (S-U).** Stacked bar chart showing protein ligand contacts plot for (S) PqsR-Taraxerone complex during the simulation of 100 ns. (T) Deviation of H-bonds contributed in interaction during 100 ns simulation. (U) Post-MD simulations intermolecular hydrogen bonding, electrostatic and hydrophobic contacts formed between Post MDS PqsR- Taraxerone complex**.** The image was generated by ligand interaction module of Schrödinger.

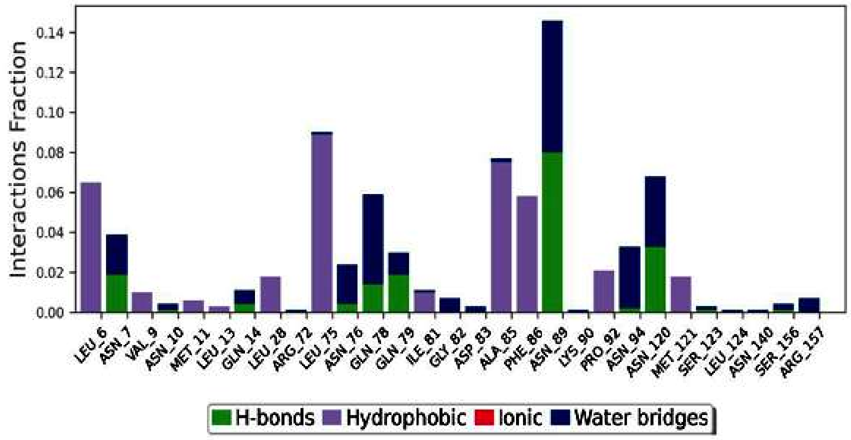


**(V)**

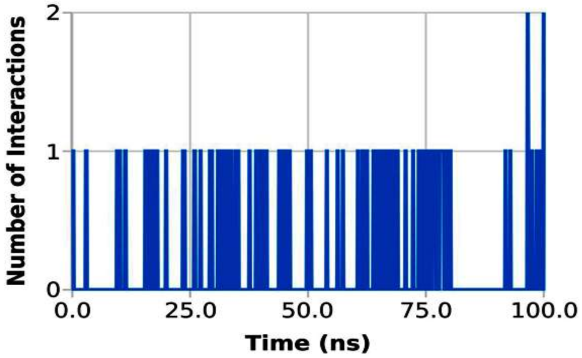


**(W)**

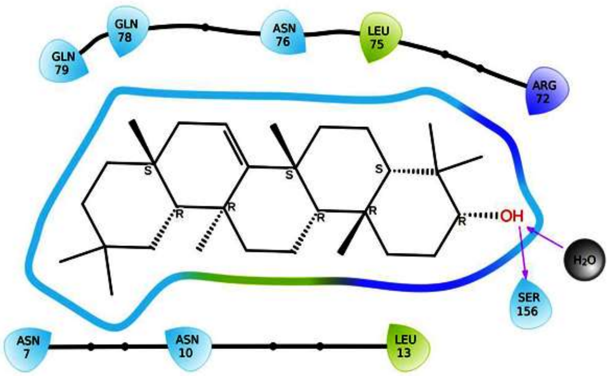


**(X)**

**Figure.: Supp1**. Stacked bar chart showing protein ligand contacts plot for (V) PqsR- Taraxerol complex during the simulation of 100 ns. (W) Deviation of H-bonds contributed in interaction during 100 ns simulation. (X) Post-MD simulations intermolecular hydrogen bonding, electrostatic and hydrophobic contacts formed between Post MDS PqsR- Taraxerol complex**.** The image was generated by ligand interaction module of Schrödinger.

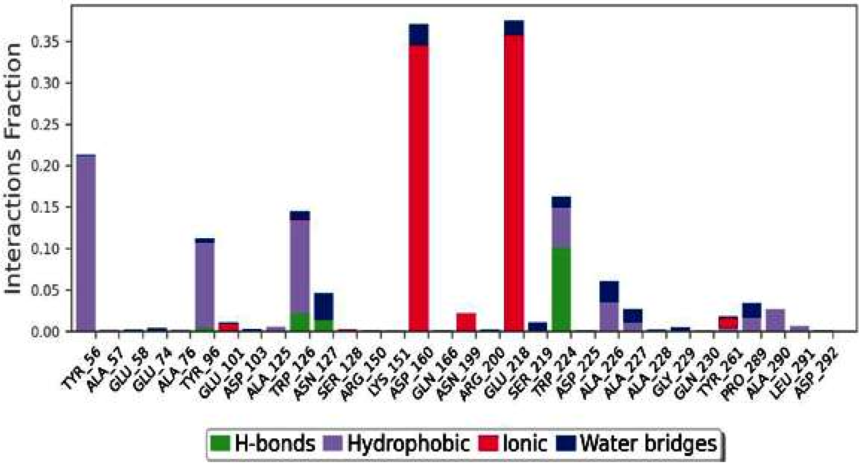


**(Y)**

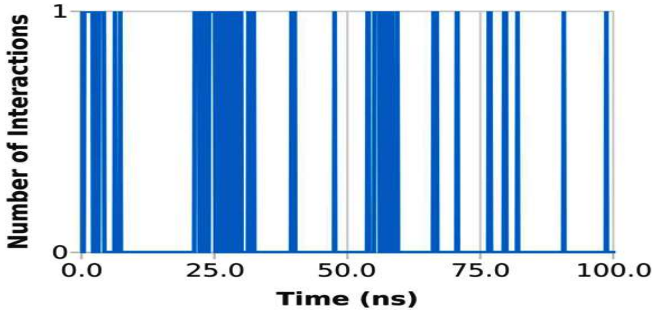


**(Z)**

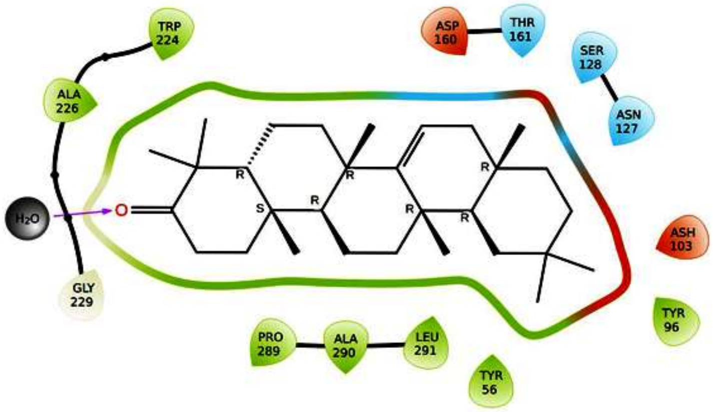


**(AA)**

**Figure S1 (Y-AA).**Stacked bar chart showing protein ligand contacts plot for (Y) pelA-Taraxerone complex during the simulation of 100 ns. (Z) Deviation of H-bonds contributed in interaction during 100 ns simulation. (AA) Post-MD simulations intermolecular hydrogen bonding, electrostatic and hydrophobic contacts formed between Post MDS pelA- Taraxerone complex**.** The image was generated by ligand interaction module of Schrödinger

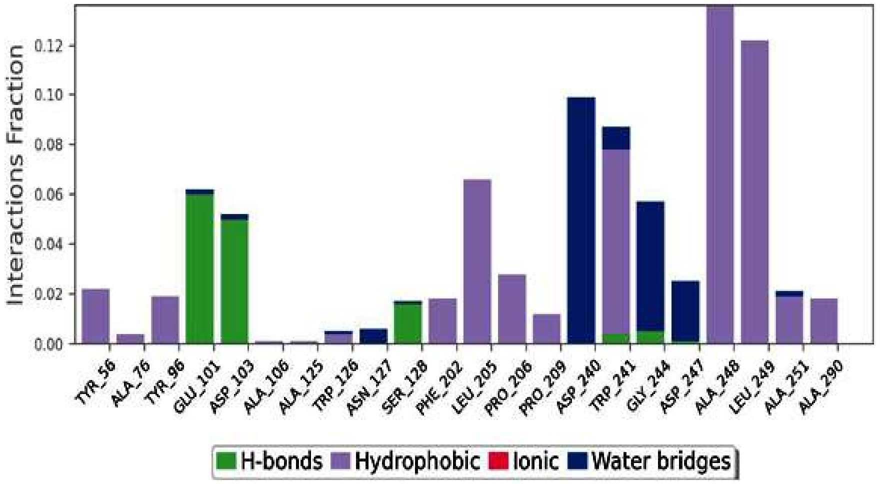


**(AB)**

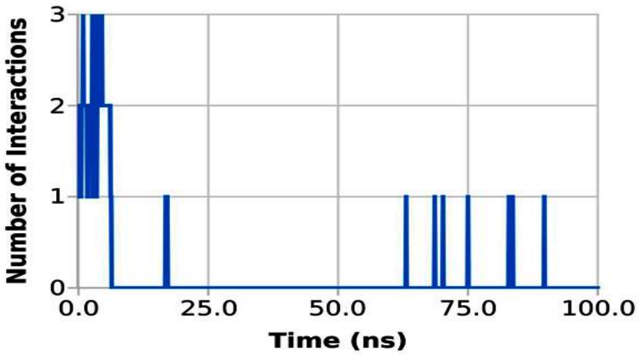


**(AC)**

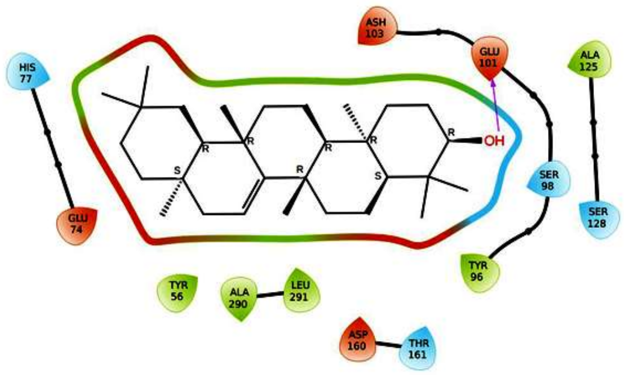


**(AD)**

**Figure S1 (AB-AD):** Stacked bar chart showing protein ligand contacts plot for (AB) PelA - Taraxerol complex during the simulation of 100 ns. (AC) Deviation of H-bonds contributed in interaction during 100 ns simulation. (AD) Post-MD simulations intermolecular hydrogen bonding, electrostatic and hydrophobic contacts formed between Post MDS PelA -Taraxerol complex**.** The image was generated by ligand interaction module of Schrödinger.

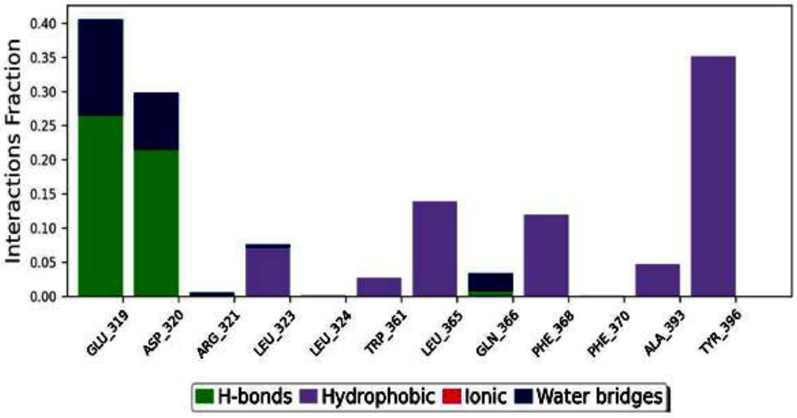


**(AE)**

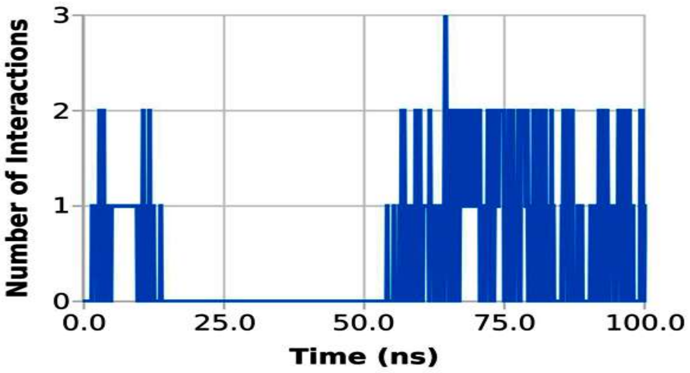


**(AF)**

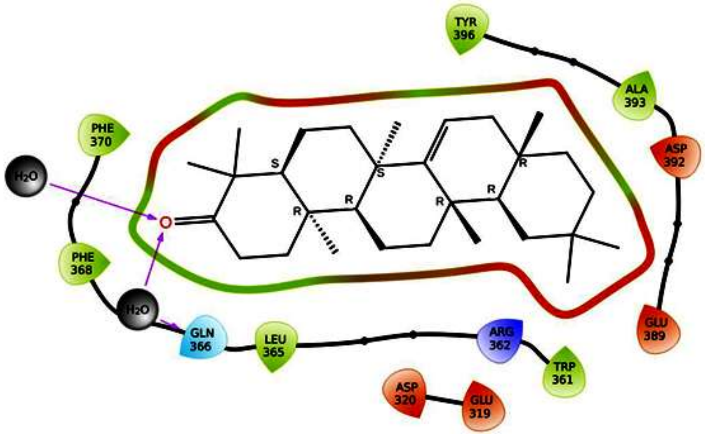


**(AG)**

**Figure S1 (AE-AG):** Stacked bar chart showing protein ligand contacts plot for (AE) PelB-Taraxerone complex during the simulation of 100 ns. (AF) Deviation of H-bonds contributed in interaction during 100 ns simulation. (AG) Post-MD simulations intermolecular hydrogen bonding, electrostatic and hydrophobic contacts formed between Post MDS PelB- Taraxerone complex**.** The image was generated by ligand interaction module of Schrödinger.

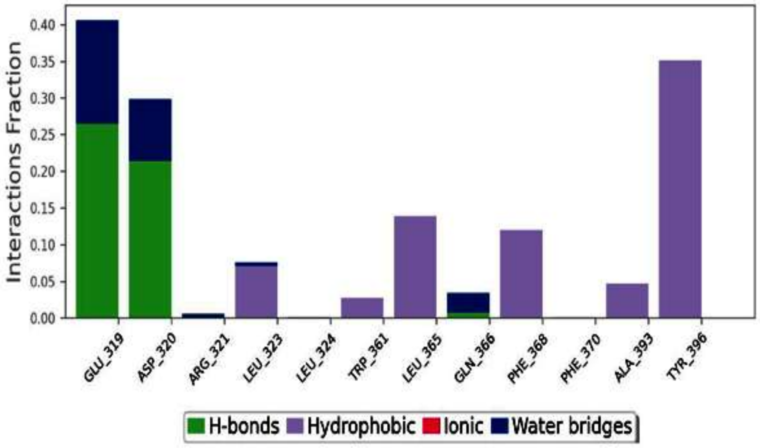


**(AH)**

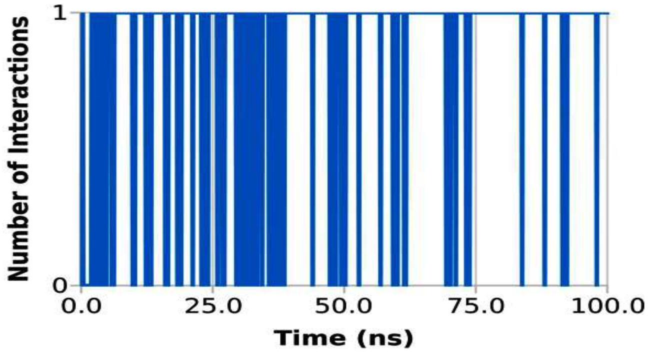


**(AI)**

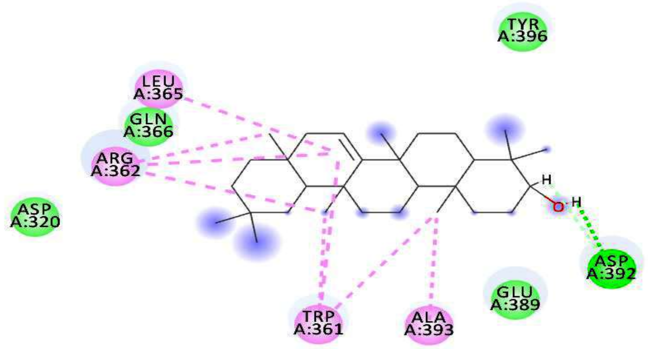


**(AJ)**

**Figure S1 (AH-AJ).: Supp1**. Stacked bar chart showing protein ligand contacts plot for (AH) PelB-Taraxerol complex during the simulation of 100 ns. (AI) Deviation of H-bonds contributed in interaction during 100 ns simulation. (AJ) Post-MD simulations intermolecular hydrogen bonding, electrostatic and hydrophobic contacts formed between Post MDS PelB-Taraxerol complex**.** The image was generated by ligand interaction module of Schrödinger.

| **Table S1:** Targets and their Binding Energies against the compounds in kcal/mol | | | | | | | | | | | | | | | | | | | | |
|---|---|---|---|---|---|---|---|---|---|---|---|---|---|---|---|---|---|---|---|---|
| **S. No** | **Phytochemical** | **Plant** | *rpoS* | *rhlR* | *rhlI* | *lasI* | *lasR* | *pqsA* | *pqSR* | *algR* | *pelA* | *pelB* | *pheA* | *pilH* | *relA* | *dinG* | *dkSA* | *spoT* | *ycgM* | *spuC* |
| 1. | 10-*O*-(*E*-cinnamoyl)-geniposidic acid | *A. marina* | -6.8 | -6.6 | -8.3 | -7 | -7.7 | -8.5 | -9 | -6.8 | -7.8 | -6.7 | -9.2 | -7 | -7.8 | -8.2 | -6.7 | -7.8 | -6.8 | -7.8 |
| 2. | 10-*O*-5-phenyl-2,4-pentadienoyl-geniposide | *A. marina* | -6.3 | -6.3 | -7.6 | -7.7 | -7.6 | -8.9 | -8.4 | -7.3 | -9 | -6.5 | -8.6 | -6.6 | -7.1 | -7.9 | -6.1 | -6.8 | -6.9 | -7.6 |
| 3. | 1-hydroxy- 8,11,13-abietatriene 12-*O*-β-xylopyranoside | *A. marina* | -6.4 | -7.3 | -7.9 | -7.5 | -7 | -8.2 | -9.2 | -6.9 | -7.5 | -8.1 | -8.8 | -6.4 | -7.5 | -9 | -7.1 | -7.5 | -6.4 | -8.5 |
| 4. | 2-(3’-3’-hydroxymethyloxiran-2’-yl-2’ methoxy-4’-Methoxymethylphenyl)-4H | *A. marina* | -5.4 | -6.4 | -7.2 | -7.5 | -6.2 | -7.8 | -8.2 | -6.3 | -7.3 | -6.4 | -7 | -5.8 | -6.5 | -8.1 | -6.2 | -7 | -6.6 | -7.3 |
| 5. | 2-[2’-2’-hydroxypropyl]-naphtha[1,2-b]furan-4,5-dione | *A. marina* | -6.2 | -7.1 | -7.9 | -6.9 | -9.9 | -8.1 | -8.3 | -7.1 | -8.2 | -7 | -8.3 | -6.3 | -6.9 | -7.8 | -6.5 | -8.2 | -6.9 | -7.2 |
| 6. | 2'-cinnamoyl-mussaenosidic acid | *A. officinalis* | -5.4 | -6.2 | -7.5 | -7.4 | -5.8 | -7.3 | -7.2 | -6.2 | -6.6 | -6 | -8.2 | -5.5 | -6.2 | -6.9 | -5.9 | -7.3 | -6.4 | -7.1 |
| 7. | 2'-*O*-(2*E*,4*E*-5-phenylpenta-2,4-dienoyl) mussaenosidic acid | *A. marina* | -6.8 | -7 | -8.3 | -7.6 | -7.9 | -9 | -8.8 | -7.4 | -7.7 | -7.1 | -7.8 | -6.3 | -7.9 | -7.7 | -7.3 | -9 | -6.4 | -8.3 |
| 8. | 3’,4’,5-trihydroxy-7-methoxyflavone | *A. marina* | -5.8 | -7 | -8 | -7.6 | -10.8 | -8.4 | -8.5 | -6.7 | -7.6 | -7.1 | -7.7 | -6.6 | -8.2 | -8 | -7.1 | -8.2 | -6.9 | -7.9 |
| 9. | 3-hydroxy-naphtha[1C-b]furan- 4,5-dione | *A. marina* | -5.7 | -6.6 | -7.8 | -6.9 | -9.5 | -7.8 | -7.4 | -6.5 | -7.9 | -6.5 | -8.2 | -5.6 | -6.5 | -7.3 | -6.6 | -7.8 | -6.7 | -6.9 |
| 10. | 4’,5,7-trihydroxyflavone | *A. marina* | -7.2 | -7 | -7.7 | -7.3 | -10.7 | -8.2 | | -6.7 | -7.2 | -6.7 | -8.4 | -6.4 | -6.9 | -7.7 | -6.8 | -7.9 | -6.7 | -7.6 |
| 11. | 4’,5-dihydroxy-3’,7-trimethoxyflavone | *A. marina* | -5.8 | -6.7 | -7.7 | -7.3 | -6.5 | -8 | -8.4 | -6.7 | -7.4 | -6.8 | -7.4 | -6.4 | -6.6 | -7.6 | -6.7 | -8.3 | -7.1 | -7.6 |
| 12. | 4’5-dihydroxy-3’-5,7-diimethoxyflavone | *A. marina* | -5.5 | -6.1 | -7.5 | -7.3 | -6.5 | -7.7 | -7.4 | -6.6 | -7 | -6.4 | -7.5 | -5.8 | -6.6 | -7.6 | -6.4 | -8.2 | -6.4 | -7.8 |
| 13. | 5-hydroxy-4; 7-dimethoxyflavone | *A. marina* | -6.1 | -7 | -7.7 | -7.8 | -10.8 | -8.5 | -8.5 | -6.4 | -7.5 | -7.3 | -7.9 | -6.5 | -8.1 | | -6.8 | -8.3 | -6.8 | -7.5 |
| 14. | 7’*S*,8’*R*-4,4’,9’-trihydroxy-3,3’,5,5’-tetramethoxy-7,8-dehydro-9-al-2,7’-cycloligan | *A. marina* | -5.4 | -5.8 | -7.2 | -6.3 | -5.8 | -6.7 | -7.3 | -5.9 | -6.5 | -5.7 | -7.8 | -5.1 | -6.3 | -6.7 | -6.1 | -6.8 | -5.8 | -6.7 |
| 15. | 7-*O*-5-phenyl-2,4-pentadienoyl-8-epiloganin | *A. marina* | -6.7 | -7.1 | -8.3 | -8.2 | -7.6 | -9.8 | -8.1 | -7.1 | -7.7 | -7.9 | -8.6 | -6.8 | -7.2 | -8.2 | -7.2 | -8.9 | -7.3 | -9.1 |
| 16. | Avicennone A | *A. marina* | -5.3 | -5.4 | -6.4 | -6.3 | -5.2 | -7 | -6.9 | -5 | -6.5 | -5.8 | -8.2 | -4.8 | -5.9 | -6 | -5.2 | -7.5 | -5.5 | -6.5 |
| 17. | Avicenol A | *A. marina* | -5.5 | -5.9 | -6.7 | -6.3 | -5.5 | -6.9 | -7.8 | -5.6 | -6.2 | -6.1 | -6.5 | -5.1 | -6.3 | -7.6 | -5.9 | -6.2 | -5.7 | -6.7 |
| 18. | Betulin | *A. marina* | -6.6 | -6.7 | -7.3 | -7.1 | -7.5 | -8.5 | -8.6 | -7.7 | -7.7 | -7.8 | -7 | -6 | -7.6 | -7 | -7.6 | -7.1 | -6.9 | -7.6 |
| 19. | Betulinaldehyde | *A. officinalis* | -6.2 | -6.9 | -7.2 | -8 | -7.6 | -9.1 | -8.6 | -7.3 | -7.9 | -7.2 | -7.1 | -6.3 | -6.9 | -8.1 | -6.5 | -7.4 | -7.4 | -8.5 |
| 20. | Betulinic Acid | *A. marina* | -6.1 | -7 | -7.2 | -7.3 | -6.9 | -8.8 | -7.5 | -7 | -7.8 | -7 | -6.8 | -6.1 | -7 | -7.7 | -6.5 | -7.5 | -7.5 | -7.6 |
| 21. | Chromen-4-one | *A. marina* | -5.1 | -7.4 | -5.8 | -5.4 | -7.6 | -6 | -5.8 | -5.2 | -6 | -5 | -6.2 | -4.6 | -4.7 | -5.4 | -4.8 | -5.8 | -5.2 | -5.4 |
| 22. | Chrysoeriol 7-O-glucoside | *A. marina* | -7.3 | -7.4 | -8.3 | -7.4 | -7.5 | -9.6 | -8.6 | -7 | -7.8 | -7.7 | -9 | -6.6 | -8.3 | -8.9 | -7.3 | -8.1 | -7.4 | -7.7 |
| 23. | ent-15- hydroxy-labda-8, 13E-dien-3-one | *A. officinalis* | -6.1 | -6.5 | -8.1 | -6.5 | -6.1 | -8.3 | -8.3 | -6.1 | -6.8 | -6.7 | -6.4 | -5.9 | -6.2 | -6.8 | -6.4 | -7.3 | -6.1 | -6.7 |

| | | | | | | | | | | | | | | | | | | | | |
|---|---|---|---|---|---|---|---|---|---|---|---|---|---|---|---|---|---|---|---|---|
| 24. | *ent*-16-hydroxy-3-oxo-13-epi-manoyl oxide | *A. officinalis* | -5.7 | -6.5 | -6.8 | -6.8 | -6.4 | -7.6 | -7.2 | -5.5 | -7 | -7.2 | -6.7 | -5.8 | -6.7 | -6.7 | -6.3 | -7.5 | -6.2 | -7.6 |
| 25. | ent-3a,15-dihydroxylabda-8,13*E*-diene, | *A. officinalis* | -5.6 | -5.9 | -7.9 | -6.5 | -5.3 | -6.5 | -6.9 | -5.5 | -6.6 | -6.9 | -7.1 | -5.6 | -5.3 | -6.8 | -5.5 | -6.9 | -6.1 | -6.5 |
| 26. | Geniposidic acid | *A. officinalis* | -6.7 | -7 | -7.3 | -7.5 | -7.9 | -9.8 | -7.2 | -7.3 | -7.9 | -6.4 | -9.2 | -5.1 | -7.4 | -9.2 | -7.5 | -7.7 | -7.2 | -8.4 |
| 27. | Isorhamnetin 3-*O*-rutinoside | *A. marina* | -6.2 | -6.7 | -9.3 | -7.5 | -7.7 | -9.8 | -9.1 | -7.4 | -7.9 | -7.4 | -8 | -6.8 | -8 | -9.8 | -7.2 | -8.6 | -7.3 | -8.5 |
| 28. | Kaempferol | *A. marina* | -5.3 | -6.9 | -7.2 | -7.2 | -6.7 | -7 | -8.1 | -6 | -6.5 | -6.5 | -5.7 | -6.8 | -6.2 | -6.9 | -6.7 | -6.8 | -6.3 | -7 |
| 29. | Lapachol | *A. marina* | -6.2 | -6.9 | -8.6 | -7.2 | -10.8 | -7 | -7.9 | -5.8 | -7.2 | -6.4 | -8 | -6 | -7 | -7.1 | -5.9 | -7.5 | -6.3 | -7.1 |
| 30. | Luteolin 7-*O*-methylether | *A. marina* | -7.7 | -6.9 | -8 | -7.4 | -11.1 | -8.5 | -8.3 | -6.5 | -7.5 | -6.8 | -9.3 | -6.8 | -6.9 | -7.8 | -7.1 | -8.1 | -7.1 | -7.9 |
| 31. | Lyoniresino | *A. marina* | -5.7 | -5.8 | -6.8 | -6.7 | -5.7 | -6.5 | -7.6 | -5.5 | -6.5 | -6 | -7.4 | -5.5 | -6.7 | -6.8 | -5.6 | -6.7 | -5.8 | -6.3 |
| 32. | Marinoids A | *A. marina* | -6.3 | -6.5 | -7.6 | -7.9 | -7.8 | -8.8 | -8.9 | -6.2 | -7.7 | -7.4 | -8.5 | -5.6 | -7.1 | -8.3 | -6.1 | -7.9 | -7.5 | -8.2 |
| 33. | Marinoids B | *A. marina* | -7.1 | -6.7 | -8.6 | -7 | -7.8 | -8.2 | -9 | -7 | -7.2 | -6.9 | -8.4 | -5.5 | -7.1 | -8.1 | -6.1 | -8.1 | -6.6 | -7.5 |
| 34. | Marinoids C | *A. marina* | -6.8 | -6.7 | -8.1 | -6.7 | -8 | -9.1 | -8.7 | -6.6 | -7 | -6.7 | -8.2 | -6.7 | -7.9 | -8.4 | -6.8 | -8 | -7 | -8.5 |
| 35. | Marinoids D | *A. marina* | -6.1 | -6.3 | -8.6 | -6.7 | -8.7 | -8.3 | -8.4 | -6.4 | -7.7 | -7.2 | -9.3 | -6.2 | -7.2 | -7.1 | -5.4 | -7.6 | -6.5 | -7.7 |
| 36. | Marinoids E | *A. marina* | -5.9 | -6.3 | -8.1 | -7.3 | -7.8 | -9 | -8.4 | -7.2 | -7 | -7.9 | -9.2 | -6.4 | -7.8 | -8.3 | -6.8 | -6.5 | -7 | -7 |
| 37. | Mussaenoside | *A. officinalis* | -5.4 | -5.8 | -7.1 | -7.1 | -5.8 | -7.7 | -7.4 | -6.3 | -6.8 | -6.1 | -8.4 | -5.6 | -6.2 | -7 | -6.2 | -6.9 | -6.3 | -6.9 |
| 38. | Naphtha[1,2-b]furan-4,5-dione | *A. marina* | -5.7 | -8.3 | -7.7 | -6.6 | -9.6 | -7.2 | -7.7 | -6.5 | -8.1 | -6.2 | -7.9 | -5.4 | -6.5 | -6.8 | -6.2 | -7.4 | -6.5 | -6.5 |
| 39. | Quercetin | *A. marina* | -6.1 | -7 | -8 | -7.4 | -11.3 | -8.3 | -8.2 | -6.4 | -7.1 | -6.5 | -8.9 | -6.6 | -7.1 | -8.1 | -6.5 | -8.1 | -6.9 | -7.3 |
| 40. | Rhizophorins A | *A. officinalis* | -4.8 | -5.2 | -6.1 | -5.2 | -4.9 | -6.1 | -6.5 | -5.1 | -5.8 | -5.4 | -8 | -4.6 | -5.9 | -5.8 | -4.9 | -5.9 | -5 | -6 |
| 41. | Rhizophorins -B | *A. officinalis* | -6.9 | -6.8 | -7 | -7 | -5.9 | -7.2 | -7.6 | -5.9 | -7.2 | -6.8 | -8.9 | -5.9 | -6.9 | -6.9 | -6.4 | -7.2 | -6.1 | -7.8 |
| 42. | Ribenone | *A. officinalis* | -6.2 | -7.7 | -7.4 | -6.8 | -6.1 | -8 | -8.3 | -6 | -7.5 | -7.1 | -6.7 | -5.6 | -6.9 | -6.9 | -6.2 | -7.2 | -6.1 | -7.3 |
| 43. | Setulin | *A. officinalis* | -5.9 | -7.7 | -7.2 | -8 | -7 | -7.9 | -10.4 | -7.3 | -8.8 | -8.1 | -9.1 | -6.6 | -7.9 | -8.7 | -7.3 | -7.7 | -6.9 | -9.5 |
| 44. | Setulinic Acid | *A. officinalis* | -6.7 | -6.9 | -7.2 | -7.3 | -6.4 | -9.2 | -7.8 | -6.9 | -7.6 | -7.9 | -7.7 | -6.4 | -7.5 | -7.6 | -6.9 | -7.8 | -6.7 | -8.8 |
| 45. | Stenocarpoquinone B | *A. marina* | -6 | -7 | -7.8 | -7.2 | -6.3 | -7.7 | -8.3 | -7.2 | -7.5 | -7 | -8.3 | -6.1 | -7 | -8 | -6.5 | -8.7 | -6.3 | -7.7 |
| 46. | Taraxerol | *A. marina* | -7.5 | -8 | -9.2 | -7.7 | -7.7 | -10.2 | -9.5 | -7.5 | -9 | -8.1 | -9.5 | -7.1 | -7.7 | -8.7 | -7 | -8.3 | -7.7 | -8.9 |
| 47. | Taraxerone | *A. marina* | -6.8 | -7.4 | -8.6 | -8.3 | -8 | -10 | -9.9 | -7.9 | -8.6 | -8.2 | -7.8 | -7 | -7.9 | -8.3 | -7.9 | -8.3 | -7.6 | -9.4 |
| 48. | Ursolic Acid | *A. marina* | -6.7 | -6.5 | -7.8 | -7.1 | -6.6 | -7.8 | -9 | -7.4 | -8.3 | -8.4 | -7.6 | -6.5 | -7.7 | -7.8 | -7.3 | -7.3 | -6.9 | -8.6 |
| 49. | B-Amyrin | *A. officinalis* | -7.1 | -7.2 | -8.6 | -7.9 | -7.7 | -9.3 | -9.5 | -6.8 | -8.5 | -8.5 | -9.4 | -7.5 | -8.3 | -8.3 | -7.6 | -7.9 | -8 | -9.8 |
